# Targeting the Mitochondrial Phenotype in Cockayne Syndrome Patient Cells: From Bioenergetic Fragility to Pharmacologic Rescue

**DOI:** 10.64898/2026.05.25.727505

**Authors:** Melis Kose, Elizabeth M. McCormick, Kelsey Keith, Cristina Remes, Suraiya Haroon, Eiko Nakamaru-Ogiso, Marni J. Falk

**Affiliations:** Mitochondrial Medicine Frontier Program, Division of Genetic and Genomic Medicine, Department of Pediatrics, The Children’s Hospital of Philadelphia, Philadelphia, PA; Department of Biomedical and Health Informatics, The Children’s Hospital of Philadelphia, Philadelphia, PA; Department of Pediatrics, University of Pennsylvania Perelman School of Medicine, Philadelphia, PA

**Keywords:** : Mitochondrial dysfunction, Complex I, Oxidative stress, antioxidants, Cockayne syndrome group B protein (CSB), *ERCC6*, Fibroblasts

## Abstract

**Background:** Cockayne syndrome (CS), primarily caused by autosomal recessive pathogenic variants in *ERCC6* (CSB) or *ERCC8* (CSA), is a transcription-coupled nucleotide excision repair disorder. CS frequently presents with features similar to primary mitochondrial disease (PMD), including leukodystrophy, lactic acidemia, and skeletal muscle mitochondrial DNA (mtDNA) depletion. How this mitochondrial phenotype arises at the cellular level, and whether it can be pharmacologically targeted, is not yet clear.

**Methods:** We characterized mtDNA content, respiratory chain (RC) protein abundance, mitochondrial biogenesis signaling pathways, and oxidative phosphorylation capacity in primary fibroblasts from two siblings with identical compound heterozygous *ERCC6* pathogenic variants (c.1526+1G>T; c.2800C>A, p.Pro934Thr) despite marked intrafamilial phenotypic divergence. A combined metabolic stress exposure (galactose, reduced glutamine, and buthionine sulfoximine, (BSO)) which reduced CS cell survival was used to screen for therapeutic leads among twenty-three candidate mitochondrial disease therapeutic compounds. Lead compounds were mechanistically validated at the level of mitochondrial superoxide, total cellular oxidative stress, glutathione, and autophagic flux.

**Results:** Patient fibroblasts exhibited several hallmarks of PMD, including reduced mtDNA content, decreased expression of complex I subunit NDUFB8, elevated expression of TOM20 with paradoxically decreased PGC1α suggestive of impaired mitophagic clearance, and decreased mitochondrial respiratory capacity. Under combined metabolic stress, ATP-levels indicative of survival in CS patient fibroblasts selectively collapsed to ∼20% of controls. Five dual-rescue compounds, defined as agents that reproducibly restored ATP-based cell survival in both patient fibroblast lines under stress, were identified, including *N*-acetylcysteine (NAC), coenzyme Q10 (CoQ10), rapamycin, taurine, and (−)-epicatechin. Mechanistic profiling resolved three functional classes of therapeutic effects in CS cells: (1) upstream mitochondrial reactive oxygen species reduction (NAC, CoQ10); (2) mTORC1 inhibition bypassing defective stress-induced autophagic induction (rapamycin); and (3) extra-mitochondrial improvement in cellular stress resilience ((-)- epicatechin, taurine).

**Conclusions:** *ERCC6*-based CSB deficiency produced a stress-sensitive and physiologically complex mitochondrial phenotype in patient fibroblasts that was pharmacologically treatable by targeting three mechanistically distinct pathways. Oxidative and broader stress buffering, autophagy modulation via mTORC1 inhibition, and enhanced cellular resilience highlight novel therapeutic opportunities to be advanced to clinical trials in CSB patients.

## Introduction

Cockayne syndrome (CS) is a rare autosomal recessive disorder that is primarily caused by pathogenic variants in *ERCC6* (CSB) or *ERCC8* (CSA), nuclear-encoded proteins that are essential for transcription-coupled nucleotide excision repair (TC-NER) (Karikkineth et al., 2017). The clinical phenotype of CS — progressive neurodegeneration, cachectic dwarfism, sensorineural hearing loss, and leukodystrophy — defines a severe, multisystem progeroid disorder with no approved disease-modifying therapy (Tiwari et al., 2021). A defining but underappreciated feature of CS is its striking phenotypic overlap with primary mitochondrial diseases (PMD): affected individuals may present with lactic acidemia, basal ganglia calcifications, and skeletal muscle mtDNA depletion — findings indistinguishable from those of classical mitochondrial disorders (Scheibye-Knudsen et al., 2012, Scheibye-Knudsen et al., 2013). Such patients have been recognized to fulfil proposed diagnostic criteria for PMD in the absence of a recognized mitochondrial disease genotype (Witters et al., 2018). This phenotypic similarity appears to reflect a mechanistic overlap rather than coincidental resemblance, yet the cellular basis of mitochondrial dysfunction in CS, and whether it can be corrected pharmacologically, remains poorly characterized (Hatch et al., 2024).

Beyond TC-NER, CSB localizes to the mitochondrial matrix, where it participates in base excision repair (BER) of oxidative lesions and interacts directly with the mitochondrial transcription and replication machinery (Aamann et al., 2010). Under conditions of nitroso-redox imbalance, loss of CSB promotes pathological accumulation of the HTRA3 serine protease, which degrades POLG — the catalytic subunit of the mitochondrial DNA polymerase —directly impairing mtDNA replication capacity (Chatre et al., 2015). CSB loss disrupts NAD+ homeostasis and impairs mitochondrial function by suppressing AMPK and ULK1 signaling. NAD+ precursor supplementation has been shown to partially rescue this defect (Okur et al., 2020). In the nucleus, CSB mediates transcription-coupled repair of DNA–protein crosslinks independently of downstream NER factors. Loss of this activity blocks transcription recovery at actively transcribed genes (Carnie et al., 2024). Together, these findings place CSB at the interface of nuclear genome surveillance and mitochondrial integrity, and help explain the PMD-like phenotype seen in patients (D’Errico et al., 2013).

Despite this mechanistic framework, key questions of mitochondrial dysfunction and potential role of therapeutics that target the mitochondrial pathophysiology of CS remain unresolved. Bioenergetic findings in CS patient-derived cells have been inconsistent. Some models show increased mitochondrial respiration, while others show broad respiratory suppression. These differences likely reflect genotype-driven variation in RC integrity, along with differences in cell type and culture conditions (Pascucci et al., 2012). The relationship between mitochondrial mass accumulation, altered mitochondrial biogenesis signaling, and organelle quality control in CSB-deficient human cells has not been resolved at the protein level. Nor has the question of whether the mitochondrial phenotype of CSB disease conditionally unmasked under defined metabolic stress been systematically addressed.

Here, we characterize the mitochondrial phenotype in primary fibroblasts from two siblings each with biparentally inherited compound heterozygous *ERCC6* variants who exhibit notable intrafamilial phenotypic divergence despite their shared genotype. A defined combined oxidative and metabolic stress paradigm to selectively reduce cell survival in CSB fibroblasts was developed, followed by screen of twenty-three candidate therapies known to target cellular pathophysiology in PMD. Five lead compounds that rescued survival in both CSB fibroblast lines were mechanistically validated at the level of mitochondrial superoxide, total cellular oxidative stress, glutathione levels, and autophagic flux.

## METHODS

### Human subjects and primary fibroblast cell lines

This study was approved by the Institutional Review Board at Children’s Hospital of Philadelphia (#08-6177, Falk, PI). Written informed consent was obtained from the legal guardians of both patients. Primary dermal fibroblasts were established from skin punch biopsies obtained from two affected siblings (Patient 1, male: Patient 2, female; both biopsies obtained at age 4 years)each carrying compound heterozygous *ERCC6* variants (c.1526+1G>T; c.2800C>A, p.Pro934Thr). Two unrelated, age- and sex-matched control fibroblast lines (CTRL 1, CTRL 2) were obtained from healthy donors having no history of photosensitivity or neurodevelopmental disease. *ERCC6* variant pathogenicity was confirmed by Sanger sequencing of exon 15 using primers flanking the c.2800C>A variant (forward: 5′-TGTGAGCGTCTTATCTGAATGG-3′; reverse: 5′-ACGAAGAGAAGGCAAAAATGC-3′). PCR was performed using Q5 High-Fidelity DNA Polymerase with 34 cycles of 30 s at 95°C, 30 s at 62°C, and 30 s at 72°C. PCR products were purified with ExoSAP-IT (ThermoFisher Scientific, #78201.1.ML) and submitted for Sanger sequencing analysis (GENEWIZ/Azenta Life Sciences).

### Cell culture

Fibroblasts were maintained in Dulbecco’s Modified Eagle Medium containing 5.5 mM glucose (DMEM 1 g/L glucose; Gibco, #11885084) supplemented with 50 µg/mL uridine and 10% fetal bovine serum (FBS; Corning, #35-010-CV) at 37°C in a humidified atmosphere of 5% CO_2_. Cultures were sub-cultured at approximately 70% confluency and used at passage thirteen or below for all experiments.

### Experimental media conditions

Three defined media conditions were used throughout this study: (1) **Basal (DMEM)**: standard growth medium, as described above. (2) **Galactose**: phenol red–free DMEM (Gibco, #21063029) supplemented to a final concentration of 10 mM D-galactose, 1 mM L-glutamine, and 10% FBS, without sodium pyruvate or uridine. Galactose substitution for glucose is a metabolic stressor that forces cells to rely on OXPHOS for ATP generation, thereby unmasking mitochondrial respiratory chain defects. (3) **Oxidative Stress**: phenol red–free DMEM supplemented to a final concentration of 10 mM D-galactose, 0.5 mM L-glutamine, 50 µM L-buthionine-sulfoximine (BSO; Sigma-Aldrich, #B2515; CAS 83730-53-4), and 10% FBS, without sodium pyruvate or uridine. BSO irreversibly inhibits γ-glutamylcysteine ligase, the rate-limiting enzyme in GSH biosynthesis, thereby depleting intracellular GSH. Combining galactose-forced OXPHOS reliance, reduced glutamine anaplerosis, and pharmacological GSH depletion creates a triple-stress challenge that reveals latent mitochondrial and redox vulnerabilities.

### Pharmacologic compounds preparation

BSO was prepared as a 100 mM stock solution in water and diluted to a final concentration of 50 µM in oxidative stress medium. Coenzyme Q10 was dissolved in DMSO. (−)-Epicatechin was dissolved in ethanol. *N*-acetylcysteine (NAC), taurine, and rapamycin were prepared as concentrated stock solutions in appropriate solvents (water for NAC and taurine; DMSO for rapamycin) and diluted into the respective media immediately before use. Vehicle controls containing matched concentrations of DMSO and/or ethanol were included in all experiments (Compound sources: L-buthionine-sulfoximine, Sigma-Aldrich, #B2515; coenzyme Q10, Acros Organics, #457950010; (−)-epicatechin, Thermo Scientific Chemicals, #J61218-F, CAS 490-46-0; *N*-acetylcysteine, Sigma-Aldrich, #A9165; taurine, Sigma-Aldrich, #T8691; rapamycin, Selleckchem, #S1039).

### ATP-based cell viability assay and candidate compound screen

Cell viability was quantified using the CellTiter-Glo Luminescent Cell Viability Assay (Promega), which measures intracellular ATP as a proxy for viable cell number. Fibroblasts were seeded into black-walled 384-well microplates at a density of 750 cells per well in 50 µL basal medium and allowed to adhere for 24 h. Medium was then exchanged to the indicated condition (basal, galactose, or oxidative stress) with or without test compounds, delivered by pin transfer from source plates (2 µL into 48 µL medium; final volume 50 µL). Each compound was tested in a minimum of four replicate wells per condition. DMSO-only controls spanning the relevant concentration range were included on each plate. At 72 h post-seeding, 25 µL CellTiter-Glo reagent was added per well. Plates were agitated for 5 min, incubated at room temperature for 15 min in the dark, and luminescence was read on a microplate reader (fixed gain 135). Values were normalized to matched vehicle-treated controls within each condition and expressed as percent viability relative to the vehicle-treated disease median.

Data analysis was performed using custom R scripts. For each well, raw luminescence values were background-subtracted (median blank well luminescence) and normalized as the percent change relative to the median luminescence of vehicle-treated disease controls on the same plate. Assay quality and dynamic range were assessed using the strictly standardized mean difference (SSMD) between positive and negative controls. Compounds were considered hits if the adjusted *q*-value was less than 0.05. All visualizations were generated using ggplot2 in R.

### Mitochondrial respiratory chain capacity analysis (Seahorse XF Cell Mito Stress Test)

Oxygen consumption rates (OCR) were measured using the Agilent Seahorse XF Pro Analyzer (Seahorse Bioscience/Agilent). Sensor cartridges from the Seahorse XFe96 Extracellular Flux Assay Kit (Agilent, #102601-100) were hydrated overnight in Seahorse XF Calibrant Solution (Agilent, #100840-000) at 37°C in a non-CO2 humidified incubator. Fibroblasts were seeded at 4 × 10⁴ cells per well in Seahorse XF96 Cell Culture Microplates (Agilent, #101085-004; included in the XFe96 Extracellular Flux Assay Kit) in basal DMEM and incubated overnight prior to the assay. On the day of the assay, Seahorse XF DMEM Medium pH 7.4 (Agilent, #103575-100) was supplemented with 10 mM glucose (Seahorse XF 1.0 M Glucose Solution; Agilent, #103577-100), 2 mM L-glutamine (Seahorse XF 200 mM Glutamine Solution; Agilent, #103579-100), and 1 mM sodium pyruvate (Seahorse XF 100 mM Pyruvate Solution; Agilent, #103578-100). Cell culture medium was exchanged for the supplemented Seahorse XF DMEM Medium and cells were incubated for 45 min at 37°C in a non-CO2 incubator to allow degassing and thermal equilibration. OCR was measured at baseline and sequentially after injection of oligomycin (1.0 µM), FCCP (2.0 µM), and rotenone/antimycin A (0.5 µM each), all supplied as part of the Seahorse XF Cell Mito Stress Test Kit (Agilent, #103015-100) and reconstituted in Seahorse XF DMEM Medium according to the manufacturer’s instructions. Each measurement phase consisted of three cycles of 3 min mix and 3 min measure (no wait). Following the assay, cells were lysed in RIPA buffer (Teknova, #R3792) supplemented with protease inhibitors, and total protein per well was quantified using the Pierce BCA Protein Assay Kit (ThermoFisher Scientific, #222325). OCR values were normalized to total protein content. Basal respiration was calculated as the OCR prior to oligomycin injection minus non-mitochondrial respiration (OCR after rotenone/antimycin A). Maximal respiration was calculated as the OCR after FCCP injection minus non-mitochondrial respiration. Spare respiratory capacity was calculated as the difference between maximal and basal respiration.

### Mitochondrial DNA copy number quantification

Relative mtDNA content was determined by SYBR Green–based quantitative PCR (qPCR). Fibroblasts were seeded in 6-well plates (patient lines at 50,000 cells per well; control lines at 30,000 cells per well) in basal DMEM and cultured for 72 hours to approximately 75% confluence prior to harvest. Genomic DNA was extracted using the Extract DNA Prep kit (QuantaBio, #95091-025) according to the manufacturer’s instructions, and samples were diluted 1:10 prior to amplification. Primers targeting the mitochondrial *MT-ND1* locus (Zhang et al., 2017) and the nuclear β2-microglobulin (*β2M*) locus (Malik et al., 2011) were used in separate reactions. Each 50 µL reaction contained 25 µL PowerUp SYBR Green Master Mix (ThermoFisher Scientific, #A25742), 1 µL each of forward and reverse primers, 5 µL template DNA, and 18 µL nuclease-free water, and was divided into three 15-µL technical replicates. Thermocycling was performed with a holding stage of 20 s at 95°C, followed by 40 cycles of 3 s at 95°C and 30 s at 60°C, and a final melt-curve analysis. Relative mtDNA content was calculated as 2^ΔCt^, where ΔCt = Ct_nuclear_ − Ct_mitochondrial_, and values were normalized to the mean of control fibroblasts processed in parallel. Three independent biological replicates were performed per cell line.

### Western immunoblot analysis

Flash-frozen cell pellets were lysed in RIPA buffer (Teknova, #R3792) supplemented with protease inhibitor cocktail and 1 mM phenylmethylsulfonyl fluoride (PMSF). Lysates were homogenized by repeated passage (five times) through a 28-gauge needle (0.5 mL insulin syringe) and incubated at 4°C for 20 min. Insoluble material was removed by centrifugation at 16,000*g* for 20 min at 4°C. Protein concentrations were determined using the Pierce BCA Protein Assay Kit (ThermoFisher Scientific, #222325). Fifty micrograms of total protein per lane were resolved on 4–15% Mini-PROTEAN TGX Precast Protein Gels (Bio-Rad, #4561084) and transferred to nitrocellulose membranes (Bio-Rad, #1620115). Membranes were blocked in 5% non-fat dry milk in TBS-T for 1 h at room temperature and incubated with primary antibodies overnight at 4°C. After washing, membranes were incubated with infrared-conjugated secondary antibodies for 1 h at room temperature. Blots were scanned on an Odyssey Infrared Imaging System (LI-COR Biosciences). Band intensities were quantified by densitometry using ImageJ (NIH).

Primary antibodies for <u>mitochondrial and biogenesis markers</u> were as follows: rabbit anti-PGC1α (Abcam, #ab191838), mouse anti-Total OXPHOS Human WB Antibody Cocktail (Abcam, #ab110411), rabbit anti-TOM20 (1:1,000; Cell Signaling Technology, #42406), and rabbit anti-citrate synthase (1:1,000; Cell Signaling Technology, #14309). Loading controls were rabbit anti-β-actin (1:1,000; Cell Signaling Technology, #4967) and mouse anti-α-tubulin (1:1,000; Sigma-Aldrich, #T5168). Secondary antibodies were IRDye 800CW goat anti-rabbit (1:10,000; LI-COR, #926-32211) and IRDye 680RD goat anti-mouse (1:10,000; LI-COR, #926-68070). Targets were normalized to β-actin or α-tubulin, as indicated in the figure legends.

<u>Autophagic activity</u> was assessed by immunoblotting for LC3-I to LC3-II conversion (LC3 lipidation) and p62/SQSTM1 protein levels under basal, galactose, and stress conditions, using the same lysis, electrophoresis, and detection protocol described above. Primary antibodies were rabbit anti-LC3B (1:1,000; Cell Signaling Technology, #2775S) and rabbit anti-SQSTM1/p62 (1:1,000; Cell Signaling Technology, #5114S). LC3 lipidation was reported as the LC3-II/LC3-I band intensity ratio; p62 levels were normalized to β-actin. To distinguish between increased autophagosome formation and impaired autophagic turnover, bafilomycin A1 (Baf A1; 100 nM) was used as a flux control. Baf A1 inhibits vacuolar H^+^-ATPase–dependent lysosomal acidification, blocking LC3-II degradation and causing its accumulation when autophagic flux is active. Accumulation of LC3-II in the presence of Baf A1 therefore confirms that the autophagic machinery is functional and that LC3 lipidation is ongoing under the experimental conditions. Rapamycin (100 nM) was tested in parallel under stress conditions to evaluate its capacity to augment LC3 lipidation. All conditions were resolved on the same gel within each experiment to enable direct comparison.

### Mitochondrial superoxide measurement (MitoSOX Red)

Mitochondrial superoxide was assessed using MitoSOX Red (Invitrogen, #M36008), a cell-permeant probe that is selectively targeted to mitochondria and exhibits fluorescence upon oxidation by superoxide (excitation/emission 510/580 nm). Fibroblasts were cultured under basal, galactose, or oxidative stress conditions (± rescue compounds) for the durations specified in the figure legends. As a positive control for mitochondrial superoxide induction, cells were treated with MitoParaquat (MitoPQ; 30 µM), a mitochondria-targeted redox cycler. On the day of the experiment, cells were first stained with 50 nM MitoTracker Deep Red FM (Invitrogen, #M22426; excitation/emission 644/665 nm) in serum-free phenol red–free medium for 30 min at 37°C to label the mitochondrial network. After washing with pre-warmed medium, cells were loaded with 500 nM MitoSOX Red prepared from a freshly dissolved 5 mM DMSO stock (13 µL DMSO per vial) in warm phenol red–free medium and incubated for 30 min at 37°C, protected from light. MitoSOX stock solutions were used within one week of preparation. Following MitoSOX staining, NucBlue Live ReadyProbes reagent (Invitrogen; 1 drop per mL of staining solution) was added as a nuclear counterstain (excitation/emission 350/461 nm). Cells were washed twice with warm PBS and imaged in phenol red–free medium on a CellInsight CX5 High-Content Screening Platform (ThermoFisher Scientific) using the following acquisition parameters: Channel 1, NucBlue (DAPI), exposure time 0.0230 s; Channel 2, MitoSOX Red (PE), exposure time 0.0978 s; Channel 3, MitoTracker Deep Red (Cy5), exposure time 0.0235 s. Acquisition settings were held constant across all conditions and experimental batches. For quantification, mitochondrial superoxide signal was extracted as MEAN_ObjectSpotTotalIntensity in Channel 2 (MitoSOX Red) and mitochondrial mass as MEAN_ObjectSpotTotalIntensity in Channel 3 (MitoTracker Deep Red), calculated as the mean per well. MitoSOX intensity was normalized to the corresponding MitoTracker Deep Red signal to control for differences in mitochondrial content between genotypes and conditions.

### Total cellular ROS measurement (CellROX Deep Red)

Total cellular reactive oxygen species (ROS) was quantified using CellROX Deep Red reagent (Invitrogen, #C10491) and flow cytometry. CellROX Deep Red is a cell-permeant probe that is non-fluorescent in a reduced state and exhibits far-red fluorescence upon oxidation by ROS (excitation/emission 644/665 nm). Fibroblasts were cultured under the indicated conditions (± compounds). On the day of the assay, CellROX Deep Red stock (supplied in DMSO; warmed to room temperature before use) was diluted into phenol red–free galactose medium to a final working concentration of 1 µM. Cells were incubated in the CellROX working solution (500 µL per well) for 30 min at 37°C, protected from light. As a positive control, fibroblasts maintained in glucose medium were treated with *tert*-butyl hydroperoxide (TBHP; 100 µM) for 1 h prior to CellROX loading. Following incubation, cells were washed once with PBS, harvested by trypsinization, and resuspended in PBS for immediate acquisition. Flow cytometry was performed on an Aurora BioSorter using the APC channel (640 nm excitation laser; 665 nm emission filter). Forward scatter/side scatter gating was used to exclude debris and doublets; single-cell populations were further refined using the R1-A (Area) parameter. Data were acquired and analysed using FCS Express 4.0 software. Unstained cells defined baseline autofluorescence. ROS levels were reported as median fluorescence intensity (MFI) and normalized to matched wild-type vehicle controls within each experimental batch. Data represent the mean ± s.d. of three independent biological replicate experiments.

### Intracellular glutathione quantification (HPLC–ECD)

Intracellular reduced glutathione (GSH) and oxidized glutathione (GSSG) were quantified by high-performance liquid chromatography with electrochemical detection (HPLC–ECD), adapted from a previously described method (Guha et al., 2021) with modifications. Frozen cell pellets were homogenized in 150 µL ice-cold water by a combination of mechanical grinding and freeze–thaw cycles. Homogenates were centrifuged at 20,000*g* for 15 min at 4°C, and the resulting supernatant was divided into aliquots for GSH (40 µL), GSSG (80 µL), and protein quantification (∼10 µL). For GSH determination, 40 µL of supernatant was deproteinized by addition of 120 µL ice-cold 1:1 (v/v) methanol:ethanol, vortexed vigorously, and kept on ice for 15 min. After centrifugation at 20,000*g* for 15 min at 4°C, the supernatant was collected and stored at −80°C until analysis. For GSSG determination, N-ethylmaleimide (NEM) was added to 80 µL of supernatant at a final concentration of 2 mM to alkylate free thiols (thereby masking GSH). Samples were vortexed for 30 s and incubated on ice for 30 min. GS–NEM conjugates and excess NEM were then removed by liquid–liquid extraction with an equal volume of ice-cold dichloromethane, followed by vortexing (15 s) and centrifugation at 1,000*g* for 10 min. The upper aqueous layer was collected, deproteinized as described above, dried under a stream of pure argon for 30 min, and stored at −80°C. Immediately before HPLC analysis, dried GSSG samples were reconstituted in water and treated with freshly prepared sodium borohydride (10 mM final concentration) for 3 min to reduce GSSG back to GSH.

Chromatographic separation was performed on a YMC-Pack ODS-A column (5 µm, 4.6 × 250 mm) with a guard column at room temperature, using a Shimadzu LC-20AD pump at a flow rate of 0.4 mL/min. The mobile phase consisted of 100 mM sodium phosphate buffer (pH 2.5) containing 75 mg/L sodium octanesulfonate and 5 mg/L EDTA. Electrochemical detection was performed with an Eicom ECD-300 detector equipped with a gold working electrode (WE-AU) and a 25 µm GS-50 gasket, operated at +400 mV against an Ag/AgCl reference electrode. Chromatograms were analysed using PowerChrom v2 software (eDAQ). GSH and GSSG concentrations were calculated from standard curves generated with freshly prepared GSH standards and normalized to total protein content of the corresponding cell homogenate.

### Statistical analyses

Data are presented as mean ± s.d. from *n* = 3 independent biological replicates unless otherwise specified. Statistical comparisons were performed using one-way ANOVA followed by Tukey’s or Dunnett’s post hoc test for multiple comparisons, or two-tailed Student’s *t*-test for pairwise comparisons, as indicated in the figure legends. *P* values are reported as: ns, not significant; \**P* < 0.05; \*\**P* < 0.01; \*\*\**P* < 0.001; \*\*\*\**P* < 0.0001. All statistical analyses were performed in GraphPad Prism (version 10).

### Data and code availability

Compound screening data and analysis scripts are available from the corresponding author upon reasonable request. All raw data underlying the figures are provided as Source Data files.

## RESULTS

### Two siblings with compound heterozygous *ERCC6* variants exhibit divergent clinical trajectories despite identical genotype

Patient 1, a 17-year-old male, exhibited a severe progressive course with regression. He achieved early motor milestones within normal limits. He walked independently at 21 months, but subsequently regressed. He was walker-dependent by age 11 years and non-ambulatory (primarily crawling) by age 15 years. Language developed to approximately 250 words intelligible to parents by age 5 years, but subsequently declining to non-verbal status by age 15 years. He had progressively decreased social engagement and was diagnosed with autism spectrum disorder at age 12 years. Growth failure was severe: at most recent evaluation (age 17 years), weight was 20 kg (Z = −14.2), height 119.1 cm (Z = −5.8), and head circumference 49.5 cm (Z = −5.2), representing 2 cm total head circumference growth from ages 2 to 17 years. Additional clinical features included cerebellar tremor and ataxia, spasticity with contractures at the elbows, knees, and ankles, motor axonal neuropathy, progressive pigmentary retinopathy, severe-to-profound bilateral sensorineural hearing loss status-post bilateral cochlear implants, gastrointestinal dysmotility with gastroesophageal reflux, and skin photosensitivity. Gastrostomy tube was required at age 7 years. Serial brain MRIs showed progressive supratentorial and infratentorial cerebral volume loss with ventriculomegaly, diffuse white matter signal abnormality, stable corpus callosum thinning, and suspected mineralization in the globus pallidus.

Patient 2, a 15-year-old female and younger sibling of patient 1, followed a relatively milder course, without neurodevelopmental regression. She walked at 24 months and at age 15 years ambulates independently with a wide-based ataxic gait. She communicates in phrases, responds to questions, and uses gestural communication. Growth failure was present but with less severe failure to thrive weight: 22.8 kg (Z = −8.4), height 115.8 cm (Z = −7.2), head circumference 47.4 cm (Z = −5.4), with 2.5 cm total head circumference growth from ages 2 to 15 years. Additional features included intention tremor with mild ataxia, mild spasticity without contractures, mild stable pigmentary retinopathy, mild-to-moderate bilateral sensorineural hearing loss (wears bilateral hearing aids), and skin photosensitivity. Gastrostomy tube was placed at 4.5 years for anticipatory feeding support and is used primarily for medications. Brain MRI at 19 months demonstrated leukodystrophy with T2-signal abnormality in the corona radiata and centrum semiovale.

### Biochemical and histopathological phenotype

Biochemical investigations revealed multiple features of mitochondrial dysfunction in both patients (Supplementary Table S1). Plasma alanine was intermittently elevated in Patient 1 (range 215.7–639.5 µmol/L over nine serial measurements between 4 years old and 17 years old; reference 89–440 µmol/L, three values above upper reference limit) and elevated in Patient 2 – age 20 months – 14 years old (251.8-603.8 µmol/L; reference 89–440 µmol/L). Anion gap was variably elevated in Patient 1 (10–20 mEq/L) and within normal limits in Patient 2 (6-13 mEq/L).

Glutathione homeostasis was concordantly impaired (Supplementary Table S1). Despite ongoing NAC supplementation (Patient 1: since 8 years of age, Patient 2: since 6 years of age; pre-treatment values not available), serial total glutathione monitoring (reference range 836–1,484 µmol/L) revealed persistently suboptimal levels in both patients, prompting iterative NAC dose escalation. In Patient 1 (ages 10–17 years), total glutathione ranged from 199 to 856.9 µmol/L, below the reference range, with NAC escalated from 10 mg/kg/day at initiation to 15 mg/kg/day (10 years) and 20 mg/kg/day (12 years). In Patient 2 (ages 8–14 years), total glutathione ranged from 314 to 1,154.4 µmol/L, with NAC escalated from 10 mg/kg/day at initiation, through 18 mg/kg/day at 8.5 years and 24 mg/kg/day at 9.5 years, to 30 mg/kg/day at 13 years following an elevated oxidized glutathione level (1.85 µmol/L; reference ≤1.78 µmol/L).

Skin biopsies obtained at age 4 years from both siblings were used to establish primary fibroblast lines. Fibroblast respiratory chain enzyme activity analysis (9 years, Baylor Genetics) demonstrated elevated rotenone-sensitive Complex I+III activity at 375% of the control mean in Patient 1 and 293% in Patient 2, after normalization to citrate synthase activity; citrate synthase activity was itself elevated in Patient 1 (148% of the control mean), suggestive of mitochondrial proliferation as an adaptive response in oxidative phosphorylation deficiency.

A vastus lateralis muscle biopsy performed in Patient 1 at age 4 years showed minimal non-specific changes with mtDNA depletion (38% of the control mean); respiratory chain enzyme activities (Baylor Genetics) were within or modestly above the laboratory reference range (Complex I 156%, Complex I+III rotenone-sensitive 131%, Complex II 132%, Complex IV 148%, citrate synthase 123% of control mean). A repeat biopsy at age 13 years revealed ragged blue fibers and subsarcolemmal mitochondrial accumulation, with partial normalization of mtDNA content (82% of the control mean) (Fig. 1d). Muscle CoQ10 quantification on the 2021 biopsy was reduced at 238 pmol/mg protein (71.4% of the control mean; reference 332.8 ± 103.0 pmol/mg). No muscle biopsy was performed in Patient 2. Biochemical and histopathological parameters for both patients are summarized in Supplementary Table S2.

**Figure 1.**
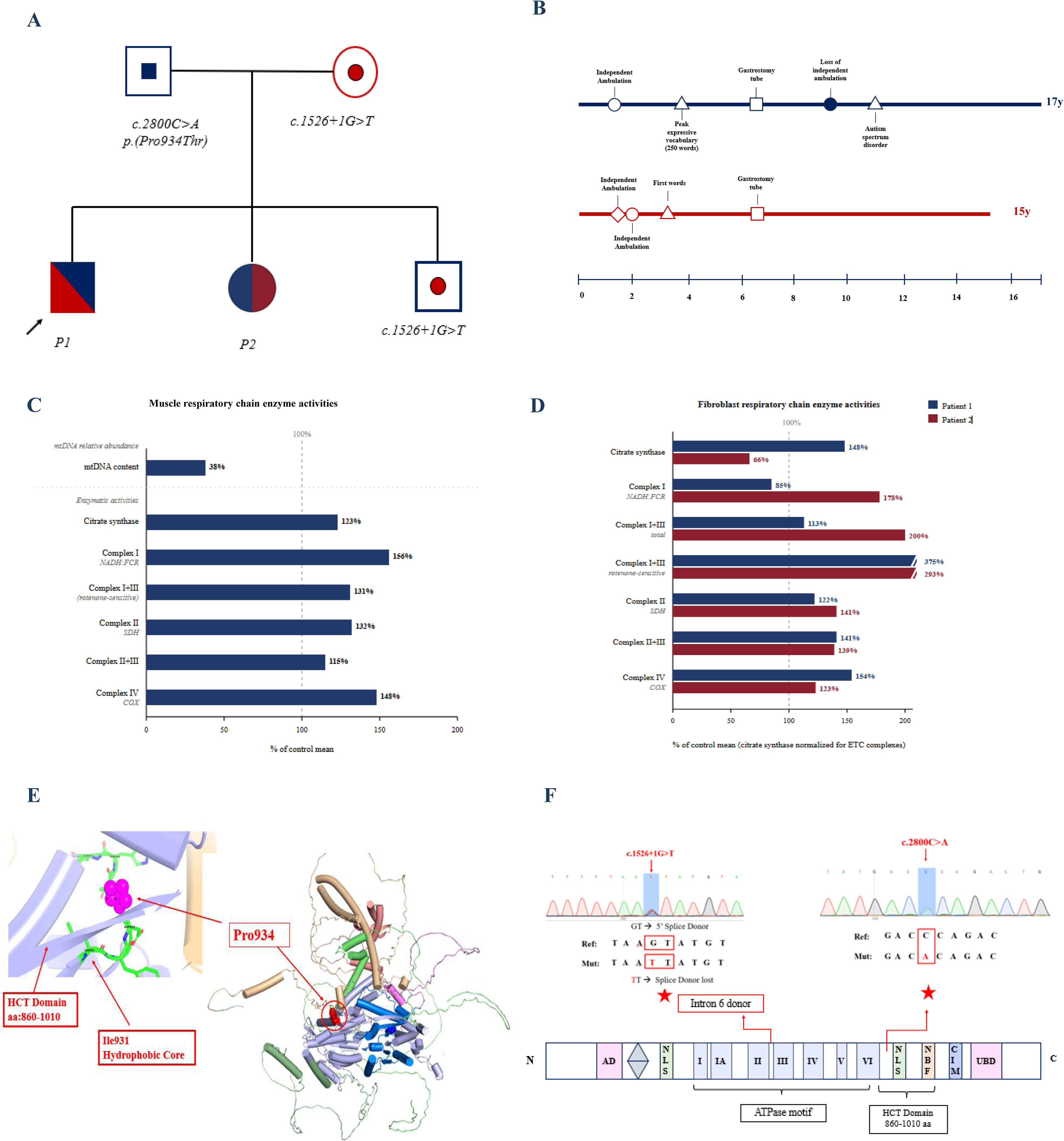
Clinical and genetic characterization of two siblings with compound heterozygous ERCC6 variants. **(a)** Pedigree of the family showing autosomal recessive inheritance. The father (I-1) carries the paternally inherited missense variant c.2800C>A (p.Pro934Thr) and the mother (I-2) carries the splice-site variant c.1526+1G>T, shown in heterozygous state. Both affected siblings (II-1, Patient 1, proband indicated by arrow; II-2, Patient 2) are compound heterozygous. The unaffected sibling (II-3) is a heterozygous carrier of c.1526+1G>T. Photographs of Patient 1 and Patient 2 are shown below the pedigree. **(b)** Longitudinal developmental timeline for Patient 1 (upper panel, severe-progressive course) and Patient 2 (lower panel, moderate-stable course), plotted from birth to current age. Color-coded horizontal tracks depict motor (Motor) and speech/language (Speech) trajectories. Annotated events include acquisition and loss of independent ambulation, peak language function, onset of regression, G-tube placement, leukodystrophy on neuroimaging, ASD diagnosis, and speech regression. Current functional status (non-ambulatory, non-verbal for Patient 1; ambulatory, phrase-level speech for Patient 2) is indicated at right. **(c)** Muscle biopsy biochemistry for Patient 1 (age 4 years). Bar graph shows respiratory chain enzyme (RCE) activities normalized to citrate synthase (CS) and expressed as percentage of the control mean. Activities measured: mtDNA content (38% of control mean), citrate synthase (CS), NADH:ferricyanide reductase-coupled Complex I (Complex I NADH:FCR), rotenone-sensitive Complex I+III (Complex I+III rot-sensitive), Complex II (SDH), Complex II+III, and Complex IV (COX). Values are indicated to the right of each bar. **(d)** Fibroblast RCE activities for Patient 1 (dark bars) and Patient 2 (light bars), normalized to citrate synthase and expressed as percentage of the control mean. Activities shown: citrate synthase, Complex I (NADH:FCR), Complex I+III total, rotenone-sensitive Complex I+III (375% for Patient 1; 293% for Patient 2, indicating selective upregulation), Complex II (SDH), Complex II+III, and Complex IV (COX). Percentage values for each patient are indicated at right. **(e)** Three-dimensional structural model of the CSB (ERCC6) protein showing the position of the p.Pro934Thr variant within the HCT domain (residues 860–1010). The HCT domain is highlighted in blue. The side chain of Pro934 (red circle) is located adjacent to the hydrophobic core residue Ile931. Structural modelling predicts that substitution of Pro934 by Thr disrupts the local backbone geometry and destabilizes the hydrophobic core, with potential impact on ATPase motif conformation. **(f)** Sanger sequencing chromatograms confirming both variants. Left: c.2800C>A (p.Pro934Thr) — reference sequence GACCAGAC versus mutant GACAGAC with C>A substitution highlighted (boxed). Right: c.1526+1G>T — reference sequence TAGTATGT versus mutant TATTTATGT with G>T transversion at the +1 position of the splice donor site, predicted to abolish canonical splicing. Bottom: schematic of the CSB/ERCC6 protein domain structure (N- to C-terminus) with positions of the NLS, ATPase helicase motifs (I, IA, II, III, IV, V, VI), NLS, CBW, CTW, and UBD domains annotated. Variant positions are indicated by red arrows.

Longitudinal assessment of leukocyte CoQ10 levels (Supplementary Table S1) revealed notable findings related to supplementation history. Although baseline leukocyte CoQ10 levels under standard ubiquinol dosing (8–16 mg/kg/day) were generally maintained at or above the upper reference limit (66–183 pmol/mg), both siblings experienced intervals of inadvertently high ubiquinol dosing (up to 300 mg/kg/day, ∼25-fold higher than standard in mitochondrial disease empiric-based clinical management). Patient 1 received supraphysiological-dose ubiquinol between approximately 8 and 10 years of age, and Patient 2 between approximately 6 and 8 years of age, during which leukocyte CoQ10 concentrations exceeded the upper control limit by 2.2- to 3.8-fold (peak 399 pmol/mg in Patient 1; peak 693 pmol/mg in Patient 2). Anecdotally, the family reported subjective clinical improvement coincident with these supraphysiological peaks and concern for clinical decline when ubiquinol or ubiquinone dosing was returned to standard empiric-based ranges (Barcelos et al., 2020).

Diagnostic confirmation in this family was protracted (Detailed diagnostic work-up is summarized in Supplementary Table S2) Initial whole exome sequencing performed at Emory Genetics Laboratory (4 years) was non-diagnostic, identifying only a paternally inherited variant of unknown significance in *SPG11* (c.4070T>C; p.L1357P) shared by both siblings. Targeted gene panels including *GJC2* (Baylor) and *GALC* sequencing, deletion, and duplication analyses (Emory Genetics Laboratory, 6 years) were similarly uninformative. Urine oligosaccharide and glycan analysis subsequently identified a reproducible abnormal peak at m/z=1449, which together with the evolving clinical phenotype prompted whole exome sequencing reanalysis. Reanalysis identified compound heterozygous *ERCC6* (NM_000124.4) variants in both siblings (Patient 1: 7 years, Patient 2: 5 years) a maternally inherited pathogenic splice-site variant c.1526+1G>T (intron 6; allele frequency 0.002% in population databases), predicted to disrupt the canonical 5′ donor splice site and to result in frameshift or nonsense-mediated decay, and a paternally inherited likely pathogenic missense variant c.2800C>A (p.Pro934Thr; exon 15; not previously reported) (Fig. 1f). Pro934 lies within the HCT domain of CSB (residues 860–1010), where the native proline imposes a backbone constraint required for hydrophobic packing with Ile931; substitution to threonine is predicted to destabilize the folded domain structure (Fig. 1e). The HCT domain mediates CSB recruitment to stalled RNA polymerase II and ubiquitin-dependent chromatin remodeling at DNA damage sites, and its structural disruption is predicted to impair both TC-NER and the mitochondrial functions of CSB. Both patients were included in a previously published patient cohort (Martin-Saavedra et al., 2022); detailed clinical features are summarized in Supplementary Table S1 (Fig. 1b, c). Both patients were included in a previously published patient cohort (Martin-Saavedra et al., 2022).

### CSB-deficient patient fibroblasts exhibited reduced mtDNA content, altered respiratory chain protein levels, and impaired mitochondrial respiratory capacity

To define the baseline mitochondrial phenotype, mtDNA content, key mitochondrial protein expression levels, and integrated respiratory capacity were quantified in patient-derived fibroblasts from both siblings compared to two unrelated control lines (Fig. 2). Quantitative PCR (*MT-ND1/β2M* ratio) analysis revealed reduced mtDNA content in both CSB-deficient fibroblast lines relative to controls, with a ∼63% reduction in Patient 1 fibroblasts and a ∼37% reduction in Patient 2 fibroblasts relative to the control mean.

**Figure 2.**
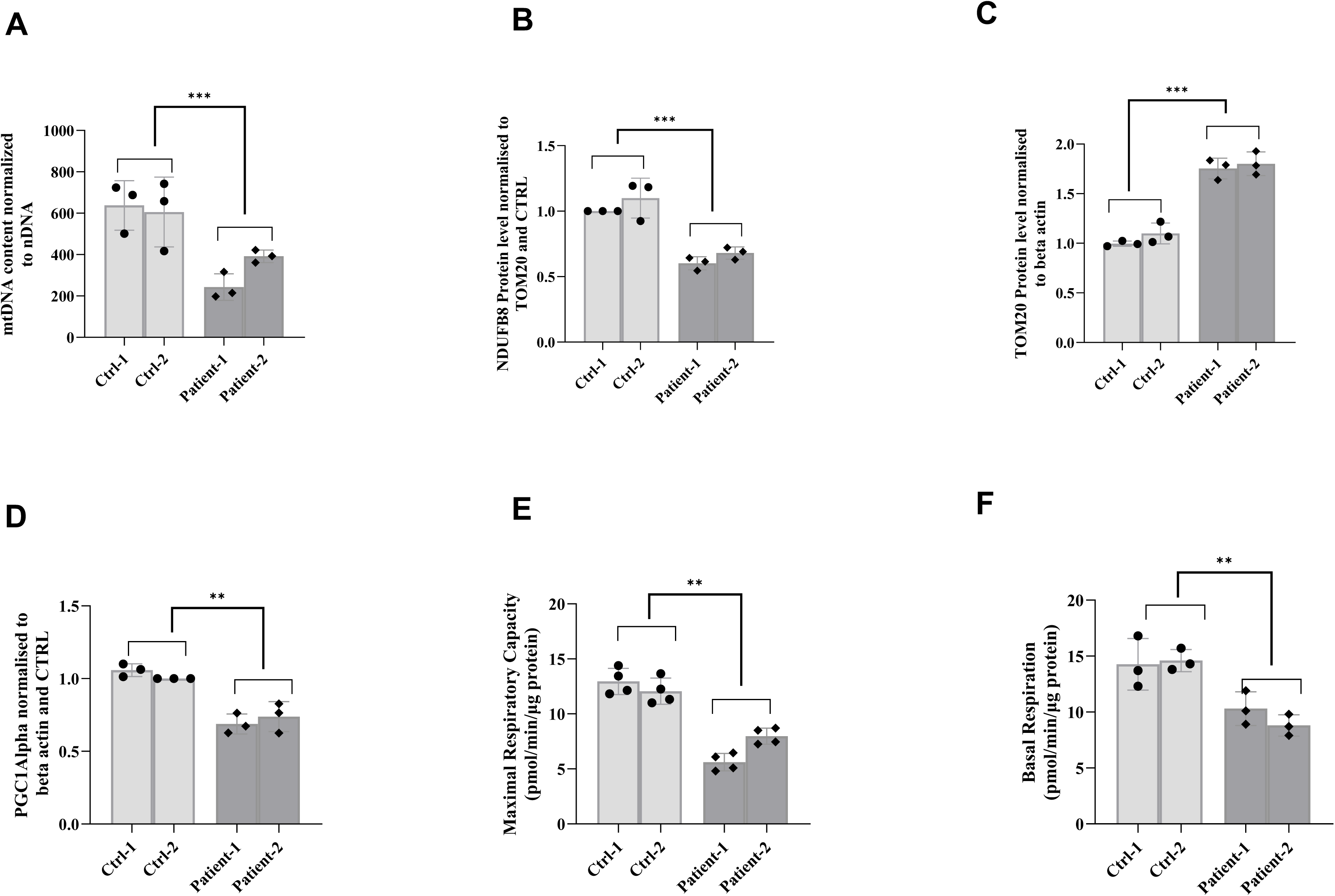
Mitochondrial phenotype of CSB-deficient patient-derived fibroblasts. **(a)** Relative mtDNA content in control (Ctrl-1, Ctrl-2) and CSB-deficient (Patient-1, Patient-2) fibroblasts, quantified by SYBR Green quantitative PCR using a mitochondrial amplicon (MT-ND1) normalized to a nuclear amplicon (β2-microglobulin; B2M). Values are expressed as mtDNA copy number normalized to nuclear DNA (nDNA). CSB-deficient fibroblasts show significantly reduced mtDNA content relative to both control lines. **(b)** NDUFB8 (mitochondrial Complex I accessory subunit) protein levels determined by immunoblotting, normalized to TOM20 protein levels and expressed relative to the control mean. CSB-deficient fibroblasts show significantly reduced NDUFB8 levels compared to controls. **(c)** TOM20 (translocase of the outer mitochondrial membrane 20; mitochondrial mass marker) protein levels determined by immunoblotting, normalized to β-actin and expressed relative to the control mean. TOM20 levels are significantly elevated in CSB-deficient fibroblasts compared to controls. **(d)** PGC1α (peroxisome proliferator-activated receptor gamma coactivator 1-alpha; master regulator of mitochondrial biogenesis) protein levels determined by immunoblotting, normalized to β-actin and expressed relative to the control mean. PGC1α levels are significantly decreased in CSB-deficient fibroblasts compared to controls. **(e)** Maximal mitochondrial respiratory capacity (oxygen consumption rate following FCCP-mediated uncoupling, minus non-mitochondrial respiration) measured by Seahorse XFe96 Cell Mito Stress Test. Values are expressed in pmol O₂/min/ µg protein. CSB-deficient fibroblasts show significantly reduced maximal respiration compared to controls. **(f)** Basal respiration (basal oxygen consumption rate (OCR)) measured under resting conditions by Seahorse XFe96 Cell Mito Stress Test, normalized to total protein content (pmol O₂/min/µg protein). CSB-deficient fibroblasts show significantly reduced basal respiration compared to controls. Data represent mean ± s.d. of n = 3 independent biological replicates per cell line; individual data points are shown. Statistical comparisons were performed by one-way ANOVA with Tukey’s post hoc test. Significance thresholds: **P < 0.01, ***P < 0.001, ; ns, not significant. *Abbreviations: CTRL, unaffected control fibroblasts; CSB, Cockayne syndrome group B protein (ERCC6); mtDNA, mitochondrial DNA; nDNA, nuclear DNA; MT-ND1, mitochondrially encoded NADH dehydrogenase 1; B2M, β2-microglobulin; NDUFB8, NADH:ubiquinone oxidoreductase subunit B8 (Complex I accessory subunit); TOM20, translocase of the outer mitochondrial membrane 20 kDa subunit; PGC1α, peroxisome proliferator-activated receptor gamma coactivator 1-alpha; OCR, oxygen consumption rate; FCCP, carbonyl cyanide-4-(trifluoromethoxy)phenylhydrazone*.

Western immunoblot analysis demonstrated decreased steady-state levels of NDUFB8 — a nuclear-encoded accessory subunit of Complex I whose stability reflects holoenzyme assembly integrity — in fibroblasts from both patient lines relative to controls (Patient 1: ∼40% reduction and Patient 2: ∼35% reduction relative to control mean). Protein levels were normalized to β-actin and expressed relative to the control mean. TOM20 protein levels, normalized to β-actin, were increased in CSB-deficient fibroblasts relative to controls in both patient lines (Patient 1: ∼1.7-fold; Patient 2: ∼1.6-fold above control mean), suggestive of mitochondrial proliferation. PGC1α protein levels, normalized to β-actin and expressed relative to the control mean, were decreased in both patient lines; this reduction reached statistical significance in Patient 1 (∼35% reduction) and showed a trend toward reduction in Patient 2 (∼28% reduction).

Seahorse XF Pro Mito Stress Test analysis demonstrated reduced basal oxygen consumption rate (OCR;∼29% reduction in Patient 1 and ∼39% in Patient 2 relative to the control mean; Fig. 2f) and reduced maximal respiratory capacity following FCCP uncoupling (∼55% reduction in Patient 1 and ∼36% in Patient 2; Fig. 2e) in CSB-deficient fibroblasts compared to controls.

Collectively, these data demonstrate that the CSB patient fibroblasts had mtDNA depletion, reduced Complex I holoenzyme assembly, and mitochondrial proliferation without increased PGC1α expression, and impaired mitochondrial respiratory chain basal and maximal capacity. Notably, consistently more severe biochemical phenotypes were present in patient 1 compared with patient 2, correlating with the more severe clinical presentation of patient 1.

### A targeted compound screen identified 5 lead compounds that rescue ATP-based survival under combined metabolic and oxidative stress

To investigate whether the mitochondrial phenotype of Cockayne syndrome is pharmacologically modifiable, we established a combined metabolic and oxidative stress condition (Fig. 3). The stress condition consisted of galactose (10 mM), reduced L-glutamine (0.5 mM), and buthionine sulfoximine (BSO; 50 µM), without sodium pyruvate or uridine. Galactose eliminates net glycolytic ATP production and forces cells to use OXPHOS; reduced glutamine restricts tricarboxylic acid (TCA) cycle anaplerosis and glutathione precursor availability; BSO inhibits γ-glutamylcysteine synthetase, depleting the glutathione pool. Basal medium consisted of DMEM with glucose (1 g/L), L-glutamine (4 mM), sodium pyruvate (1 mM), and uridine (1 mM). Cell survival was quantified by ATP-based luminescence analysis after 72 hours in stress media relative to basal media.

**Figure 3.**
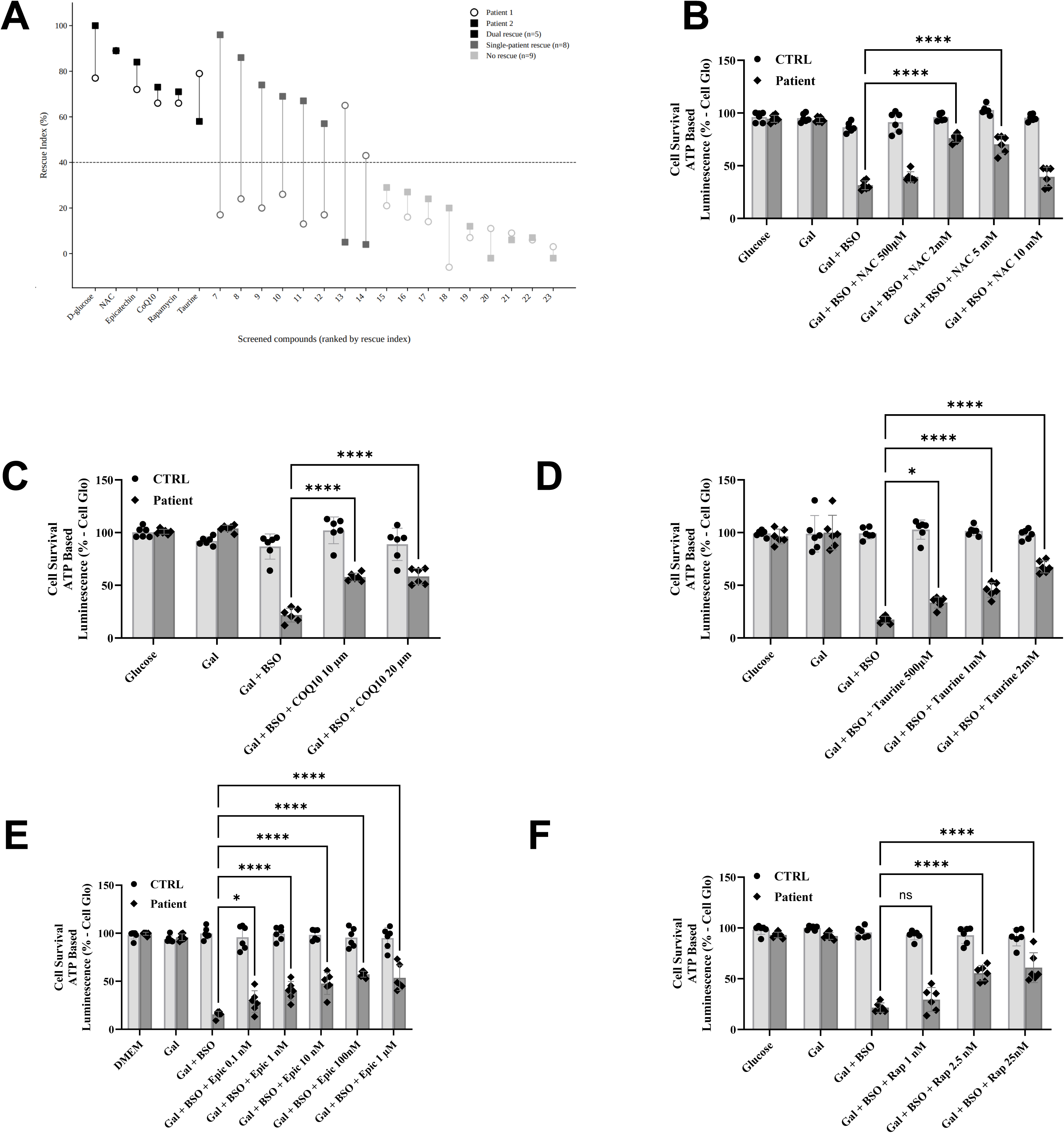
Targeted compound screening identifies five agents that rescue ATP-based cell survival selectively in CSB-deficient fibroblasts under combined metabolic stress. **(a)** Scatter plot showing rescue index (%) for each compound in Patient 1 (x-axis) versus Patient 2 (y-axis) fibroblasts under the stress condition (galactose 10 mM, L-glutamine 0.5 mM, BSO 50 µM, without sodium pyruvate or uridine). Rescue index is calculated as: RI (%) = (Viabᵉcompound+BSO+Gal − ViabᵉBSO+Gal) / (Viabᵉunstressed − ViabᵉBSO+Gal) × 100. The dashed line indicates the 40% rescue threshold. Compounds achieving RI ≥ 40% in both patient lines (dual-rescue hits) are shown in black; named dual-rescue compounds are labelled. Non-hit compounds are shown in grey (identities embargoed; Cmp 1–17). D-glucose (positive control) is shown in dark grey, positioned leftmost on the x-axis. Compounds with patient-selective rescue are labelled by patient line. **b–f,** Dose–response validation of the five dual-rescue compounds — **(b)** NAC, **(c)** taurine, **(d)** coenzyme Q10 (CoQ10), **(e)** (−)-epicatechin, and **(g)** rapamycin — in CTRL and CSB-deficient (Patient-1, Patient-2) fibroblasts. Cells were cultured under basal DMEM (5.5 mM D-glucose, 4 mM L-glutamine, 1 mM sodium pyruvate, 1 mM uridine), galactose alone (10 mM galactose, 0.5 mM L-glutamine), or stress (10 mM galactose, 0.5 mM L-glutamine, 50 µM buthionine sulfoximine, BSO) at the indicated compound concentrations. Total intracellular ATP was quantified by CellTiter-Glo luminescence assay and expressed as percent of CTRL under basal DMEM. CTRL data represent pooled measurements from two unrelated control fibroblast lines (Ctrl-1, Ctrl-2). Patient data represents pooled measurements from two sibling fibroblast lines carrying identical compound heterozygous ERCC6 variants (Patient-1, Patient-2). Patient-1 and Patient-2 showed concordant responses across all 32 conditions tested (Pearson r = 0.97; no condition reached statistical significance after Benjamini–Hochberg correction); data were therefore pooled for visualization. Per-cell-line raw values, per-condition statistical tests, and a concordance scatter plot visualizing these data are provided in Source Data Figure 3. Bars represent mean ± s.d. Statistical comparisons were performed by one-way ANOVA with Tukey’s post hoc test. \**P* < 0.05, \*\*\*\**P* < 0.0001; ns, not significant. *Abbreviations: BSO, buthionine sulfoximine (inhibitor of γ-glutamylcysteine synthetase); Gal, galactose; DMEM, Dulbecco’s Modified Eagle Medium; NAC, N-acetylcysteine; CoQ10, coenzyme Q10 (ubiquinone); Epi, (−)-epicatechin; Rap, rapamycin; CTRL, unaffected control fibroblasts; ATP, adenosine triphosphate; RI, rescue index; Cmp, compound)*.

Under basal DMEM conditions containing glucose, CSB-deficient fibroblasts exhibited cell survival and ATP levels comparable to that of control fibroblasts. In contrast, the combined stress condition caused a sharp reduction in ATP-based survival in CSB-deficient fibroblasts, falling to approximately 20% of basal DMEM levels, while control fibroblasts were largely unaffected (Fig. 3). This hypersensitivity to combined metabolic and oxidative stress provided a screening platform to test pharmacological rescue.

A targeted panel of 23 compounds representing distinct mechanistic classes that our research group has previously found to have therapeutic benefit in preclinical models of PMD — including antioxidants, electron transport chain cofactors, autophagy modulators, and metabolic intermediates (Falk, 2021, Guha et al., 2021, McCormack et al., 2015, Peng et al., 2015)— was screened in parallel across both CSB-deficient patient 1 and patient 2 fibroblast lines (Fig. 3a, b). Compounds were co-administered with the stress condition media (galactose (10 mM), reduced L-glutamine (0.5 mM), and buthionine sulfoximine (BSO; 50 µM), without sodium pyruvate or uridine) for 72 h. Screening outcomes were categorized as: rescue in both patient fibroblast lines, rescue in one patient fibroblast line only, or no rescue in either line. A subset of compounds demonstrated patient-line-specific effects, improving ATP-based survival in only one of the two fibroblast lines. However, 5 compounds that reproducibly restored ATP-based cell survival in both CSB-deficient patient fibroblast lines were selected for downstream validation on the basis of this dual-line rescue activity: N-acetylcysteine (NAC), taurine, coenzyme Q10 (CoQ10), (−)-epicatechin, and rapamycin. Compound identities for the remaining screened compounds are designated by numerical code (Cmp 1–18).

To confirm reproducibility and assess dose dependence, the 5 compounds that rescued both cell lines were subsequently evaluated for dose–response effects in control and CSB-deficient fibroblasts cultured under basal DMEM, galactose alone, and combined metabolic and oxidative stress conditions. Each compound produced a significant, dose-dependent increase in ATP-based cell survival in CSB-deficient fibroblasts specifically under combined stress conditions, with no significant effect under basal DMEM conditions (Fig. 3b-f).

Because NAC is a direct cysteine donor that replenishes the GSH pool targeted by BSO, we specifically asked whether GSH precursor supplementation retains efficacy when initiated after stress-induced GSH depletion has already occurred (Aldini et al., 2018, Atkuri et al., 2007).To assess whether NAC retained rescue activity when initiated only after stress onset, NAC was added 48 hours after BSO exposure. Importantly, ATP-based survival at 96 hours post-BSO (48 hours after NAC addition) was significantly higher than at the 48-hour stressed baseline (Supplementary Fig. S1), indicating that the therapeutic rescue window in CSB-deficient cells extends beyond the initial period of acute GSH depletion.

### Mitochondrial superoxide levels are constitutively elevated in CSB-deficient fibroblasts and differentially modulated by the 5 lead candidate therapies

To determine whether the 5 lead compounds modify mitochondrial oxidative redox status, we quantified mitochondrial superoxide relative levels using the MitoSOX probe on a CellInsight CX5 high-content imaging platform. MitoSOX fluorescence mean intensity per well was normalized to MitoTracker Deep Red (MtrDR) fluorescence mean intensity per well to control for mitochondrial mass differences across conditions and cell lines. MitoParaquat (MitoPQ), a mitochondria-targeted redox cycler that selectively generates superoxide at the Complex I flavin site (Robb et al., 2015), was used as a positive control to establish the maximal MitoSOX signal (Fig. 4a). Under basal DMEM conditions, CSB-deficient fibroblasts exhibited significantly higher MitoSOX/MtrDR signal than control fibroblasts, indicating constitutive elevation of mitochondrial superoxide (Fig. 4). Culturing cells in galactose alone did not significantly alter this signal. However, the combined metabolic and oxidative stress condition produced a statistically significant further increase in MitoSOX/MtrDR intensity selectively in CSB-deficient fibroblasts.

**Figure 4.**
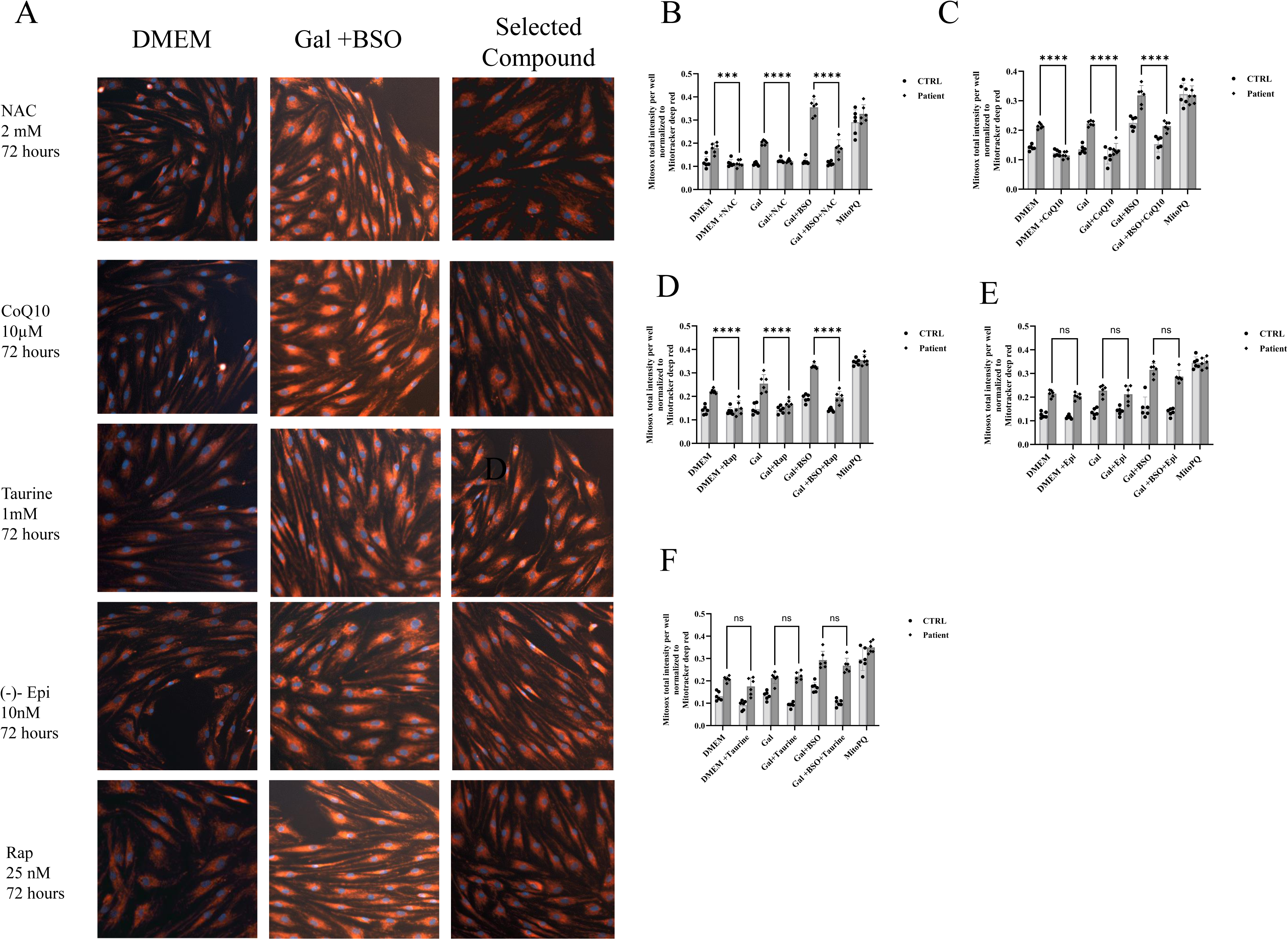
Mitochondrial superoxide is constitutively elevated in CSB-deficient fibroblasts and is differentially modulated by the five rescue compounds. **(a)** Representative fluorescence micrographs acquired on a CellInsight CX5 High Content Screening platform (Thermo Fisher Scientific) showing mitochondrial superoxide (MitoSOX Red; excitation 510 nm, emission 580 nm) and mitochondrial mass (MitoTracker Deep Red, MtrDR; excitation 644 nm, emission 665 nm) in control and CSB-deficient (Patient-1) fibroblasts. Nuclei were counterstained with NucBlue Live ReadyProbes Reagent (DAPI equivalent; excitation 360 nm, emission 460 nm) for spatial normalization. Cells were imaged under the following conditions: basal DMEM (glucose 1 g/L, L-glutamine 4 mM, sodium pyruvate 1 mM, uridine 1 mM); combined stress (galactose 10 mM, L-glutamine 0.5 mM, BSO 50 µM, without sodium pyruvate or uridine); and stress in the presence of each indicated rescue compound. MitoParaquat (MitoPQ; mitochondria-targeted redox cycler that generates superoxide selectively at the Complex I flavin site) was used as a positive control to define the maximal inducible MitoSOX signal. A comprehensive panel of micrographs including galactose-only and all compound conditions is provided in Supplementary Figure 4. **b–f,** Quantification of normalized mitochondrial superoxide in control (CTRL) and CSB-deficient (Patient) fibroblasts under basal DMEM, galactose alone, galactose + BSO (stress), and stress + compound conditions, as indicated. MitoSOX total fluorescence intensity per well was normalized to MitoTracker Deep Red total fluorescence intensity per well (MitoSOX/MtrDR ratio) to account for mitochondrial mass differences between genotypes. MitoPQ defines the upper reference limit of the assay dynamic range. Compound panels: **(b)** NAC (2 Mm- 72 hours) — significantly reduced MitoSOX/MtrDR signal in CSB-deficient fibroblasts under both basal and stress conditions; **(c)** CoQ10 (10 µM – 72 hours) — significantly reduced MitoSOX/MtrDR signal under both basal and stress conditions; **(d)** rapamycin (25 nM – 72 hours) — significantly reduced MitoSOX/MtrDR signal under both basal and stress conditions; **(e)** (−)-epicatechin (10 nM – 72 hours) — did not significantly alter MitoSOX/MtrDR signal under any condition tested; **(f)** taurine (1mM-72 hours) — did not significantly alter MitoSOX/MtrDR signal under basal, galactose, or stress conditions. Effects were specific to the CSB-deficient genotype; no significant alterations were detected in control fibroblasts across all treatments. Data represent mean ± s.d. of n = 3 independent biological replicates per cell line; individual data points are shown. Statistical comparisons were performed by one-way ANOVA with Tukey’s post hoc test. Significance thresholds: ***P < 0.001, ****P < 0.0001; ns, not significant. Fluorescence quantification was performed using CellInsight CX5 HCS software (Thermo Fisher Scientific); a minimum of 200 cells per well were analysed per condition. *Abbreviations: MitoSOX, MitoSOX Red mitochondrial superoxide indicator; MtrDR, MitoTracker Deep Red FM (mitochondrial mass marker); MitoPQ, MitoParaquat; BSO, buthionine sulfoximine; Gal, galactose; NAC, N-acetylcysteine; CoQ10, coenzyme Q10; Epi, (−)-epicatechin; Rap, rapamycin; DMEM, Dulbecco’s Modified Eagle Medium; CTRL, unaffected control fibroblasts; HCS, high-content screening*.

Candidate therapy mechanistic effects on mitochondrial superoxide burden were not uniform across the 5 lead compounds that rescued cell survival in CSB-deficient fibroblasts (Fig. 4b–f). NAC (Fig. 4b), CoQ10 (Fig. 4c), and rapamycin (Fig. 4f) significantly reduced MitoSOX/MtrDR in CSB-deficient fibroblasts under both basal DMEM and combined stress conditions (Fig. 4b). In contrast, neither taurine (Fig. 4d) nor (−)-Epicatechin (Fig. 4e) significantly altered MitoSOX/MtrDR under basal, galactose, or combined stress conditions (Fig. 4d). Of note, the observed reductions in mitochondrial superoxide burden with NAC, CoQ10, and rapamycin were specific to the CSB-deficient genotype, as no significant alterations were observed in control fibroblasts across any treatment or condition. Overall, these analyses demonstrated that mitochondrial superoxide levels are constitutively elevated in CSB-deficient fibroblasts and only rescued by 3 of the 5 lead candidate therapies found to rescue cell survival. Glutathione redox dynamics confirmed stress paradigm validity.

Intracellular glutathione levels (GSH, reduced; GSSG, oxidized; and the GSH/GSSG redox ratio) were measured in Patient 1 fibroblasts and compared to Control 1 fibroblasts under basal DMEM, galactose, and combined stress conditions (Fig. 5a–c). Baseline GSH concentrations and GSH/GSSG ratios did not differ significantly between control and CSB-deficient fibroblasts under basal or galactose-only conditions. BSO-containing stress media produced near-complete depletion of reduced GSH and collapse of the GSH/GSSG ratio in both genotypes, confirming effective γ-glutamylcysteine synthetase inhibition. NAC (2 mM, 72 hours) treatment showed a directional trend toward GSH replenishment under BSO-containing conditions (Fig. 5a). Interestingly, CoQ10 (25 µM, 72 h) treatment significantly restored the GSH/GSSG ratio despite not serving as a direct glutathione precursor. Thus, only two of the 5 candidate therapies, namely NAC and CoQ10, rescued glutathione levels under combined metabolic and oxidative stress in CSB-deficient fibroblasts.

**Figure 5.**
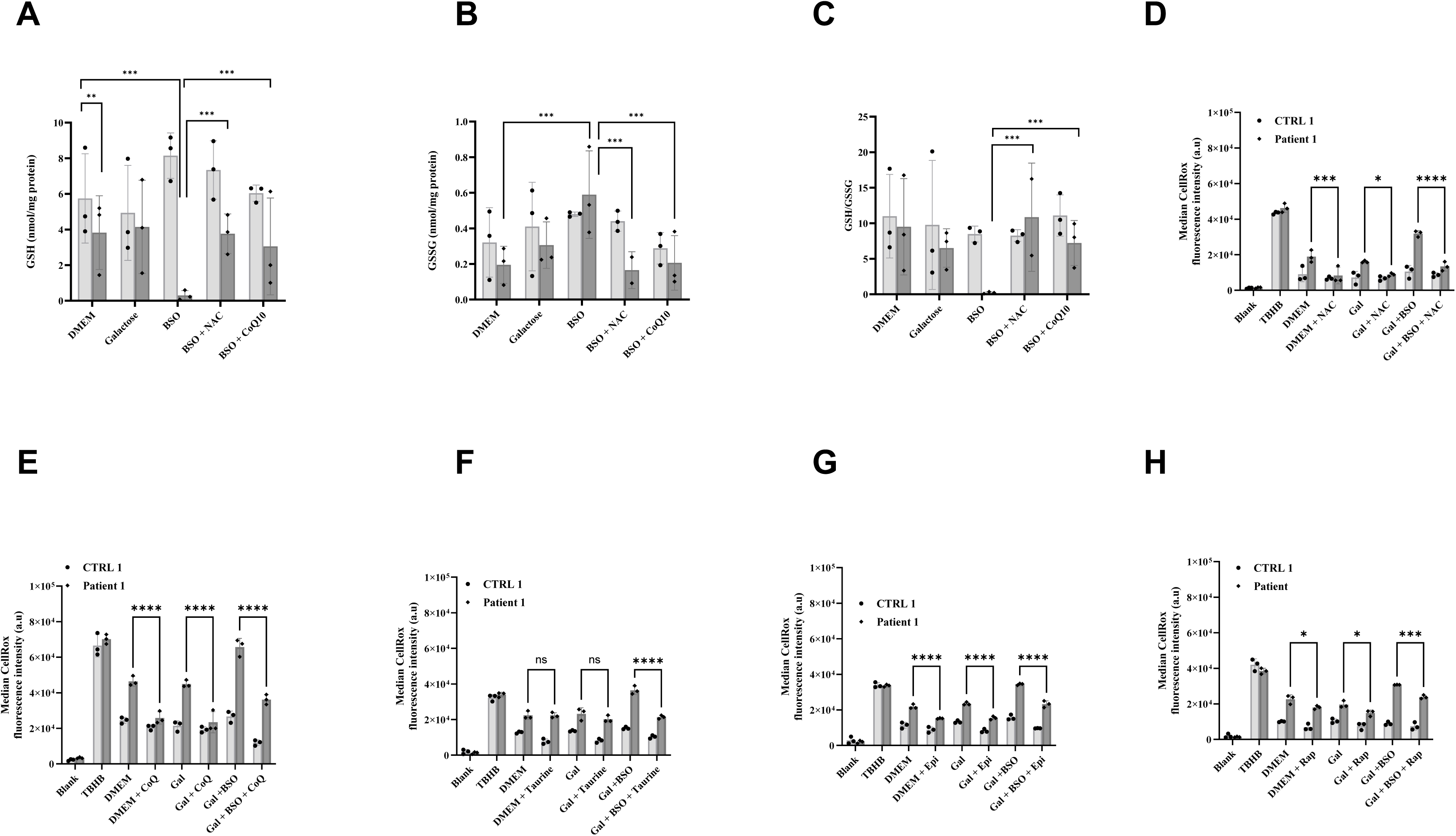
Biochemical validation of glutathione dynamics and flow cytometric quantification of total cellular ROS under stress and rescue conditions. **a–c,** Intracellular glutathione indices quantified in one unaffected control line (Control-1) and the more severely affected CSB-deficient line (Patient-1) under the following conditions: basal DMEM (glucose 1 g/L, L-glutamine 4 mM, sodium pyruvate 1 mM, uridine 1 mM); galactose alone (10 mM, L-glutamine 4 mM); BSO stress (galactose 10 mM, L-glutamine 0.5 mM, BSO 50 µM, without sodium pyruvate or uridine); BSO + NAC; and BSO + CoQ10. Glutathione species were measured using the GSH/GSSG-Glo Luminescent Assay (Promega) according to the manufacturer’s instructions. **(a)** Reduced glutathione (GSH; nmol/mg protein). **(b)** Oxidised glutathione (GSSG; nmol/mg protein). **(c)** GSH/GSSG ratio. BSO treatment induces a significant reduction in GSH levels and collapse of the GSH/GSSG ratio in Control-1 and Patient-1, confirming effective γ-glutamylcysteine synthetase blockade. NAC supplementation shows a trend toward GSH restoration under BSO conditions. CoQ10 significantly restores the GSH/GSSG ratio despite not functioning as a direct glutathione precursor. **d–h,** Total cellular ROS quantified by flow cytometry in one unaffected control line (Control-1) and the more severely affected CSB-deficient line (Patient-1) using CellROX Deep Red Reagent (Thermo Fisher Scientific; excitation 644 nm, emission 665 nm; APC channel, 640 nm excitation laser). Fibroblasts were incubated with CellROX Deep Red (5 µM, 30 min, 37°C) prior to analysis on a BD LSRFortessa flow cytometer. Single-cell populations were identified and gated on the R1-A (area) scatter parameter in FCS Express software (De Novo Software). Data are expressed as median fluorescence intensity (MFI) normalized to the Control-1 DMEM condition. Tert-butyl hydroperoxide (TBHP; 200 µM, 30 min) was included as a positive control to define the maximal inducible oxidative shift in the APC channel; unstained cells were used to establish background autofluorescence baseline. Under basal DMEM, Patient-1 fibroblasts exhibit constitutively elevated CellROX MFI relative to Control-1. Compound response profiles under basal DMEM, galactose, and galactose + BSO (stress) conditions: **(d)** NAC — significantly reduced CellROX MFI across all three conditions; **(e)** CoQ10 — significantly reduced CellROX MFI across all three conditions; **(f)** taurine — no significant reduction under basal DMEM or galactose; significantly reduced CellROX MFI under galactose + BSO stress; **(g)** (−)-epicatechin — significantly reduced CellROX MFI across all three conditions including basal DMEM; (h) rapamycin — significantly reduced CellROX MFI across all three conditions Data represent mean ± s.d. of n = 3 independent biological replicates per cell line; individual data points are shown. Statistical comparisons were performed by one-way ANOVA with Tukey’s post hoc test. Significance thresholds: *P < 0.05, **P < 0.01, ***P < 0.001, ****P < 0.0001; ns, not significant. *Abb:GSH, reduced glutathione; GSSG, oxidised glutathione; BSO, buthionine sulfoximine (γ-glutamylcysteine synthetase inhibitor); NAC, N-acetylcysteine; CoQ10, coenzyme Q10; Epi, (−)-epicatechin; Rap, rapamycin; Gal, galactose; DMEM, Dulbecco’s Modified Eagle Medium; CellROX, CellROX Deep Red Reagent (oxidative stress indicator); MFI, median fluorescence intensity; TBHP, tert-butyl hydroperoxide (positive control oxidant); APC, allophycocyanin; ROS, reactive oxygen species; CTRL, unaffected control fibroblasts*.

### Total cellular ROS analysis revealed 3 distinct compound response profiles

Total cellular reactive oxygen species (ROS) were assessed by flow cytometry analysis using CellROX Deep Red, a cell-permeant dye whose fluorescence increases upon oxidation by ROS. Patient 1 fibroblasts exhibited constitutive elevation of CellROX MFI relative to controls under basal DMEM conditions (2-fold). Compound responses could be stratified into three distinct profiles: [1] NAC, CoQ10, and rapamycin significantly reduced total cellular ROS under combined stress by 58 % (p < 0.0001), 45% (p < 0.0001), and 23% (p < 0.001), respectively (Fig. 5d, e, h). [2] (−)-Epicatechin significantly reduced total cellular ROS by 33% across all conditions (p < 0.0001; Fig. 5g), but had yielded no measurable reduction of mitochondrial superoxide burden (Fig. 4e, 5g); and [3] Taurine reduced total cellular ROS significantly only under BSO-containing combined stress conditions (41%; p < 0.0001) without effect under basal DMEM or galactose-only conditions (Fig. 5f).–h), and yielded no rescue of mitochondrial superoxide burden under any condition.

Collectively, these studies demonstrate that both mitochondrial superoxide burden and total cellular oxidative stress are significantly rescued under conditions of combined metabolic and oxidative stress by NAC, CoQ10, or rapamycin. By contrast, (-)-epicatechin and taurine did not rescue mitochondrial superoxide levels and had differential effects on total cellular ROS, where (-)-epicatechin was beneficial under basal and stressed conditions while taurine showed significant benefit only under conditions of oxidative stress.

#### CSB-deficient fibroblasts had impaired stress-induced autophagosome formation, which was augmented by rapamycin

To directly evaluate the autophagic response given the observed ability of rapamycin (a known modulator of mTORC2-mediated autophagy (Kim and Guan, 2015)) to reduce CSB-deficient cell line survival, LC3-I/LC3-II conversion and p62 (SQSTM1) levels were assessed by western immunoblot analysis in Patient 1 fibroblasts alongside a control line under basal DMEM, galactose, and combined stress conditions (Fig. 6a). Under basal DMEM conditions, LC3-II/LC3-I ratios and p62 levels normalized to β-actin were comparable between control and CSB-deficient fibroblasts, with no significant genotype-dependent difference (Fig. 6b, c). However, under combined metabolic stress (galactose + BSO + reduced glutamine), CSB-deficient fibroblasts showed a significant decrease in LC3-II/LC3-I ratio relative to stressed controls, along with failure to clear p62, reflecting impaired autophagy induction.

**Figure 6.**
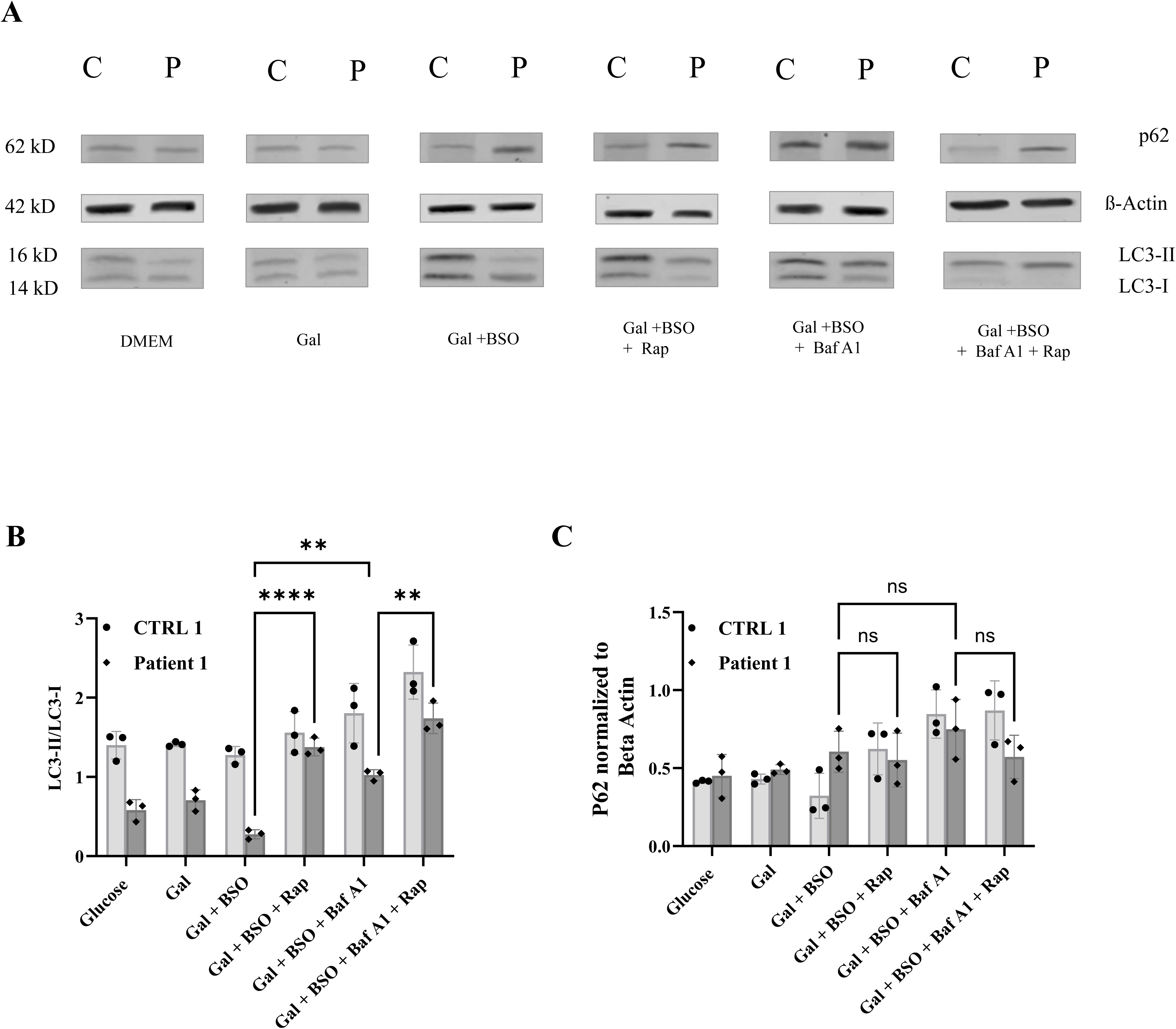
CSB-deficient fibroblasts exhibit impaired stress-induced LC3 lipidation that is rescued by rapamycin. **(a)** Representative immunoblots for p62 (SQSTM1; 62 kDa), β-actin (42 kDa), LC3-I (16 kDa), and LC3-II (14 kDa) in one unaffected control line (Control-1; C) and the more severely affected CSB-deficient line (Patient-1; P) under the following conditions (left to right): basal glucose/DMEM; galactose (10 mM, L-glutamine 1 mM, without sodium pyruvate and uridine); galactose + BSO stress (galactose 10 mM, L-glutamine 0.5 mM, BSO 50 µM, without sodium pyruvate or uridine); stress + rapamycin (Rap; 100 nM); stress + bafilomycin A1 (Baf A1; 25 nM); and stress + Baf A1 + rapamycin. Molecular weight markers are indicated. **(b)** Quantification of the LC3-II/LC3-I ratio by densitometry from immunoblots shown in panel a. Under basal glucose/DMEM, LC3-II/LC3-I ratios are comparable between control and CSB-deficient fibroblasts. Under galactose + BSO stress, the LC3-II/LC3-I ratio is significantly lower in CSB-deficient fibroblasts than in stressed controls (****P < 0.0001). Rapamycin significantly increases the LC3-II/LC3-I ratio under stress in both control (****P < 0.0001) and CSB-deficient fibroblasts (**P < 0.01). Bafilomycin A1 results in LC3-II accumulation in both genotypes, confirming intact autophagic machinery and lysosomal degradation capacity. Combined Baf A1 + rapamycin maintains elevated LC3-II/LC3-I levels at or above Baf A1 alone (**P < 0.01 versus stress alone). **(c)** Quantification of p62 levels normalized to β-actin by densitometry. Under basal glucose/DMEM and galactose-alone conditions, p62 levels are comparable between genotypes (ns). Under galactose + BSO stress, p62 levels are not significantly different between control and patient fibroblasts. Rapamycin does not significantly alter p62 levels under stress conditions (ns). Bafilomycin A1 results in significant p62 accumulation in both genotypes (ns between genotypes under Baf A1). Combined Baf A1 + rapamycin maintains p62 accumulation at levels comparable to or greater than Baf A1 alone Data represent mean ± s.d. of n = 3 independent biological replicates per cell line; individual data points are shown. Statistical comparisons were performed by one-way ANOVA with Tukey’s post hoc test. Significance thresholds: *P < 0.05, **P < 0.01, ***P < 0.001, ****P < 0.0001; ns, not significant. *LC3, microtubule-associated protein 1A/1B-light chain 3 (autophagosome marker); LC3-I, cytosolic (unlipidated) form; LC3-II, phosphatidylethanolamine-conjugated (lipidated, autophagosome-associated) form; p62, sequestosome-1 (SQSTM1; autophagy cargo receptor); Baf A1, bafilomycin A1 (vacuolar H⁺-ATPase inhibitor; blocks autophagosome–lysosome fusion); Rap, rapamycin (mTORC1 inhibitor; induces autophagy); BSO, buthionine sulfoximine; Gal, galactose; DMEM, Dulbecco’s Modified Eagle Medium; C, control fibroblasts; P, CSB-deficient patient fibroblasts; mTORC1, mechanistic target of rapamycin complex I*.

To assess whether the reduced LC3-II/LC3-I ratio reflected impaired autophagosome formation or accelerated LC3-II turnover, cells were treated with bafilomycin A1 (Baf A1), a specific inhibitor of the vacuolar H⁺-ATPase that blocks autophagosome–lysosome fusion and prevents LC3-II degradation. Baf A1 treatment led to clear LC3-II and p62 accumulation in both control and CSB-deficient fibroblasts (Fig. 6b, c), indicating that the autophagic flux machinery itself is intact (Yamamoto et al., 1998). The reduced LC3-II/LC3-I ratio under stress conditions therefore reflects impaired autophagosome formation rather than accelerated turnover. Rapamycin (25Nm, 72 hours) treatment under combined stress conditions significantly increased the LC3-II/LC3-I ratio in both control and patient 1 fibroblasts (5-fold; p < 0.0001), and combined Baf A1 plus rapamycin maintained LC3-II and p62 levels at or above those observed with Baf A1 alone in both genotypes. These data are consistent with CSB-deficient having impaired autophagosome formation and autophagic flux, which is rescued by treatment with rapamycin.

## DISCUSSION

Two siblings with Cockayne syndrome caused by identical compound heterozygous *ERCC6* variants presented with clinical features indistinguishable from PMD (Gorman et al., 2016) including gross motor developmental delay, progressive neurodevelopmental regression, ataxia, leukodystrophy with basal ganglia involvement, growth failure, pigmentary retinopathy, sensorineural hearing loss, elevated blood GDF-15 and plasma alanine, muscle biopsy showing mtDNA depletion (below the diagnostic cut-off of 50% of control mean), and elevated citrate synthase activity with ragged red fibers consistent with compensatory mitochondrial proliferation (Labory et al., 2020). These findings satisfy previously proposed diagnostic criteria for PMD in the absence of a known genetic etiology (Witters et al., 2018). At the cellular level, patient-derived fibroblasts recapitulated key biochemical hallmarks of mitochondrial dysfunction — mtDNA depletion, reduced respiratory chain complex I abundance, suppressed respiratory chain capacity, and impaired organelle quality control at the level of autophagy — that collectively provide a mechanistic basis for their clinical phenotype that overlapped with that of clinical mitochondrial disease (Karikkineth et al., 2017, Scheibye-Knudsen et al., 2013). Of therapeutic relevance, this cellular phenotype was pharmacologically modifiable — five compounds, acting through distinct pathways, reproducibly restored viability in both patient fibroblast lines under combined metabolic and oxidative stress.

Cellular mtDNA content was reduced in fibroblasts by ∼63% in Patient 1 and by ∼37% in Patient 2 relative to controls, in line with their clinical severity gradient and consistent with CSB’s established role in mitochondrial genome maintenance through BER at the mtDNA level (Aamann et al., 2010, Kamenisch et al., 2010). Further, steady-state NDUFB8 levels — an accessory subunit whose abundance reflects the stability of assembled Complex I holoenzyme — were reduced in both patient lines (Stroud et al., 2016). This cellular finding contrasts with elevated rotenone-sensitive Complex I+III activity in fibroblast respiratory chain enzyme activity assays (375% and 293% of control mean after citrate synthase normalization). This contrast might be explained by a distinction between maximal catalytic throughput of the residual enzyme, which may also be upregulated as a result of compensation, and the total quantity of correctly assembled complex, which is reduced at the protein level (McGregor et al., 2023, Stroud et al., 2016). Indeed, similar findings are common in Leber’s Hereditary Optic Neuropathy (LHON) mitochondrial complex I disease models (Giordano et al., 2014, Lin et al., 2012). The elevated citrate synthase activity in the clinical muscle biopsy of patient 1 is consistent with a parallel compensatory mitochondrial proliferation response occurring at the tissue level (Larsson and Oldfors, 2001).

The increase in mitochondrial mass in CSB-deficient fibroblasts reflects impaired mitochondrial clearance rather than reduced biogenesis. TOM20, a mitochondrial outer membrane protein, was elevated in both CSB-deficient patient fibroblast lines, consistent with with the elevated citrate synthase activity in clinical biopsy material. However, PGC1α that is the master transcriptional activator of the mitochondrial biogenesis program (Halling and Pilegaard, 2020) was significantly reduced in Patient 1 (p < 0.01) and trended toward reduction in Patient 2. Increased mitochondrial mass is normally driven by PGC1α-dependent biogenesis. The opposite pattern observed here — elevated TOM20 alongside reduced PGC1α — indicates that mitochondrial accumulation in CSB-deficient cells reflects insufficient organelle clearance rather than increased biogenesis (Palikaras and Tavernarakis, 2014). Indeed, CSB is required for the recruitment of autophagy factors to damaged mitochondria, and impaired mitophagic flux in CSB-deficient models has been reported to result in the progressive retention of dysfunctional organelles (Scheibye-Knudsen et al., 2012, Scheibye-Knudsen et al., 2013).

The CSB-deficient fibroblasts showed evidence of bioenergetic suppression. Basal and maximal OCR capacity were both reduced in CSB-deficient fibroblasts, indicating that the respiratory chain operates below its innate capacity despite the presence of elevated mitochondrial mass. This respiratory suppression phenotype has been reported in some CS cellular models (Pascucci et al., 2012) but contrasts with a hyper-active respiration phenotype that was observed in CSB-null mouse embryonic fibroblasts (Scheibye-Knudsen et al., 2012). The combination of reduced mtDNA content, decreased Complex I NDUFB8 abundance, and suppressed OCR despite elevated mitochondrial mass defines a specific bioenergetic state in which mitochondrial quantity is maintained while functional output is disproportionately reduced — a profile consistent with impaired organelle quality control (Palikaras and Tavernarakis, 2014). This pattern reflects loss of the direct mitochondrial maintenance and quality control functions of CSB (Aamann et al., 2010) rather than an epiphenomenal mitochondrial response to the nuclear transcription-repair defect (Viscomi and Zeviani, 2020).

We identified a combined stress condition that selectively revealed the bioenergetic vulnerability of CSB-deficient cells. Under basal media conditions, CSB-deficient fibroblasts-maintained ATP-based viability comparable to controls, indicating that the mitochondrial phenotype is partially compensated at rest. Under combined metabolic and oxidative stress (galactose, reduced glutamine, and BSO) cell viability decreased to ∼20% of basal rates in CSB-deficient patient lines while control lines remained unaffected. Each stressor adds a distinct challenge: galactose forces OXPHOS-dependent ATP synthesis by eliminating glycolytic rescue; reduced glutamine restricts both TCA cycle anaplerosis and GSH precursor supply; BSO pharmacologically dismantles the GSH pool by blocking γ-glutamylcysteine ligase (Griffith and Meister, 1979). The selective decrease in CSB-deficient cells is consistent with a baseline near the limit of redox tolerance, which becomes critical only when antioxidant capacity and metabolic flexibility are both reduced.

Five compounds with distinct mechanisms reproducibly rescue CSB-deficient fibroblast cell viability. Upon screening of 23 compounds previously identified in our research studies to rescue preclinical models of primary mitochondrial disease, we identified five compounds that also reproducibly restored ATP-based viability in both CSB-deficient patient fibroblast lines. These 5 lead therapeutic compounds included NAC, CoQ10, rapamycin, taurine, and (−)-epicatechin. Several additional compounds improved survival in only one of the two lines, which may reflect intrafamilial divergence of other genetic modifying factors. Follow-up profiling of mitochondrial superoxide levels, total cellular ROS levels, and glutathione indices stratified the 5 lead compounds into 3 mechanistic categories according to their effects across compartmental and contextual oxidative stress readouts: Mitochondrial superoxide burden and total cellular oxidative stress were significantly rescued under conditions of combined metabolic and oxidative stress by NAC, CoQ10, or rapamycin. Neither (-)-epicatechin nor taurine rescued mitochondrial superoxide levels but had differential effects on total cellular ROS, where (-)-epicatechin was beneficial under basal and stressed conditions while taurine showed significant benefit only under conditions of oxidative stress.

NAC and CoQ10 showed mitochondrial oxidative redox restoration in CSB-deficient patient fibroblasts. Both NAC and CoQ10 significantly reduced mitochondrial superoxide production under basal and stress conditions, reduced total cellular ROS across all conditions, and restored the GSH/GSSG ratio under BSO stress. NAC functions as a cysteine donor that replenishes intracellular GSH, bypassing the BSO-mediated enzymatic block at γ-glutamylcysteine ligase (Griffith and Meister, 1979). Under BSO-induced GSH depletion the mitochondrial GSH pool, which is imported from the cytosol via the inner membrane carrier SLC25A39, becomes rate-limiting for matrix hydrogen peroxide detoxification. The reduction in MitoSOX fluorescence by NAC is consistent with restored GSH-dependent antioxidant function (Wang et al., 2021).

Unlike NAC, CoQ10 acts through a complementary dual mechanism. First, as an obligate two-electron carrier between Complexes I/II and Complex III, CoQ10 may reduce electron leak at the FMN site of partially assembled Complex I (Hirst and Roessler, 2016), consistent with the NDUFB8 reduction observed in patient cells. Second, in its reduced ubiquinol form CoQ10 functions as a lipophilic membrane antioxidant (Quinzii et al., 2012). The significant restoration of the GSH/GSSG ratio by CoQ10, without its acting as a direct glutathione precursor, indicates that its primary effect is a reduction in the oxidant burden that drives GSH consumption — a mechanistically distinct route to redox restoration as compared with NAC. Notably, NAC retained rescue activity even when added 48 hours after stress onset, after cell viability had already dropped, indicating that the therapeutic window for GSH precursor supplementation extends beyond the point of acute depletion (Supplementary Fig. S1). The effectiveness of post-stress NAC suggests that sustained mitochondrial oxidant generation—rather than the immediate loss of the GSH pool—drives cell death in this model. This finding has direct implications for therapeutic timing in patients. Both CSB-deficient patients accidentally received supraphysiological ubiquinol doses, which raised leukocyte CoQ10 levels several-fold above the control range. Parents reported improved clinical function during this period; although anecdotal, this observation is consistent with our cellular finding that higher tissue CoQ10 may offer therapeutic benefit in Cockayne syndrome (Aristizabal-Henao et al., 2026, Subbiah et al., 2025).

Rapamycin therapy both reduced oxidative stress and induced autophagy by bypassing the CSB-dependent induction defect. Rapamycin reduced mitochondrial superoxide and total cellular ROS levels across diverse conditions and significantly increased the LC3-II/LC3-I conversion ratio in patient and control fibroblasts under stress. Under metabolic stress, CSB-deficient fibroblasts failed to upregulate LC3 lipidation relative to stressed control cells, despite having intact autophagic flux machinery as was confirmed by demonstrating bafilomycin A1 accumulation of LC3-II in both genotypes. This finding suggests the defect relates to impaired stress-responsive induction rather than turnover, and is consistent with the requirement for CSB in recruiting autophagy machinery to damaged organelles (Scheibye-Knudsen et al., 2012, Scheibye-Knudsen et al., 2013). Rapamycin bypasses this induction defect through direct mTORC1 inhibition (Sarkar et al., 2009), restoring LC3-II accumulation and clearing the autophagy substrate p62 (Kim and Guan, 2015). The indirect mechanism by which rapamycin lowers mitochondrial superoxide via selective removal of the most depolarized, ROS-generating organelles rather than by direct antioxidant activity is consistent with this quality control model. Whether the affected cargo is mitochondria-selective (mitophagy) or involves broader macroautophagy will need to be addressed with mitophagy-specific reporters such as mito-Keima or mito-QC (Kajitani et al., 2025). The autophagy induction defect and its rescue by rapamycin in CSB-deficient cells are similar to our previously published findings in PMD models, where impaired LC3B lipidation and dysregulated p62 clearance also accompanied respiratory chain dysfunction; in those studies, rapamycin along with lithium chloride and 3-methyladenine restored cell viability and preserved respiratory capacity across genetic and pharmacological RC-deficient models (Peng et al., 2015). The shared autophagy phenotype and its response to pharmacological correction across CS and PMD point to a common downstream vulnerability in the mitochondrial stress response that can be targeted by mTORC1-directed therapy. (−)-Epicatechin provided extra-mitochondrial, compartment-specific redox rescue in CSB-deficient cells. While (−)-epicatechin reproducibly restored cell viability in both patient lines and reduced total cellular ROS across all basal and stress conditions, it did not significantly reduce mitochondrial superoxide levels under any condition. This subcellular compartmental dissociation demonstrates that cell viability rescue under the stress condition does not require direct mitochondrial superoxide reduction, but can also be achieved by enhancing redox capacity in the cytosolic compartment. Epicatechin activates the AMPK–PGC1α axis and induces NRF2-mediated antioxidant gene expression (McDonald et al., 2021, Taub et al., 2012) — pathways that are both suppressed in CSB-deficient patient 1 fibroblasts (as demonstrated by reduced PGC1α expression and constitutive CellROX elevation) and have been shown to be consistently dysregulated across Cockayne syndrome transcriptomes (Kajitani et al., 2025). Epicatechin also disrupts the NRF2–KEAP1 interaction, inducing phase II detoxification enzymes (NQO1, HO-1) that operate predominantly in the cytosolic and microsomal compartments (Chiou et al., 2016). Induction of these antioxidant enzymes (SOD2, GPx1, NQO1, HO-1) lowers total cellular ROS (CellROX) without affecting mitochondrial superoxide levels (MitoSOX), which is consistent with the observed profile.

Taurine treatment in CSB-deficient cells demonstrated a context-dependent tolerance mechanism orthogonal to direct ROS scavenging. Taurine did not reduce mitochondrial superoxide or total cellular ROS under basal or galactose-only stress conditions. However, it rescued viability under BSO-containing oxidative stress and lowered total cellular ROS specifically when GSH was depleted. This strict context-dependence, namely inactive against the constitutive redox burden but protective when the GSH-dependent antioxidant barrier is acutely dismantled, suggests taurine functions as a tolerance factor upon exposure to acute oxidative stress rather than as a classical antioxidant. Two non-mutually exclusive mechanisms are most relevant to the CS context. First, taurine is incorporated as 5-taurinomethyluridine (τm⁵U) into mitochondrial tRNAᴸᵉᵘ(UUR) by the MTO1–GTPBP3 complex, a modification required for efficient UUG codon decoding during mitochondrial translation, including of MT-ND6, a complex I subunit (Kopajtich et al., 2014, Suzuki et al., 2002). The recently characterized inner mitochondrial membrane taurine transporter SLC6A6 establishes a direct molecular pathway through which exogenous taurine can sustain this modification (Li et al., 2026);–in CSB-deficient cells with reduced Complex I subunit levels, as maintaining translational output under acute oxidative stress may be critical for preserving residual respiratory function. Second, taurine functions as a membrane-stabilizing osmolyte (Schaffer et al., 2010) when the GSH-dependent antioxidant barrier is removed by BSO, since membrane integrity may become an independent rate-limiting determinant of survival (Singh et al., 2023). Interestingly, taurine has obtained regulatory agency approval in Japan as a preventative therapy for strokes in one classical mitochondrial disease, namely mitochondrial encephalopathy, lactic acidosis, and stroke-like episodes (MELAS) syndrome (Ohsawa et al., 2019). The normal plasma taurine status of both siblings (Supplementary Table S1) indicates that the cellular rescue observed here does not reflect correction of a systemic taurine deficiency state, but rather aligns with the recent demonstration that intramitochondrial taurine availability—rather than extracellular taurine concentration—is the critical determinant of mitochondrial translational output (Li et al., 2026). This distinction supports evaluation of taurine supplementation as a prospective intervention in CS cohorts, specifically targeting the intramitochondrial pool that SLC6A6 is required to replenish.

Intriguingly, the two CSB-deficient siblings reported here showed substantial intrafamilial phenotypic divergence and patient-specific pharmacological responses. Despite sharing an identical *ERCC6* compound heterozygous genotype, the two siblings show markedly divergent clinical trajectories. At last evaluation in their mid-teens, Patient 1 had sustained pronounced neurodevelopmental regression and was nonambulatory and nonverbal, while Patient 2 had no reported regression episodes and remained ambulatory, communicative, and socially interactive. Intrafamilial variability of this magnitude has been documented in CS as attributed to secondary genetic or epigenetic modifiers (Calmels et al., 2018). In our compound screen, a subset of compounds improved survival in only one patient fibroblast line, reflecting a pharmacological response heterogeneity that mirrored the cellular heterogeneity between the two lines (greater mtDNA depletion and PGC1α reduction in patient 1) and may well reflect the same biological divergence that underlies the clinical difference

Notably, during the intervals of inadvertent supraphysiological-dose ubiquinol exposure beyond that standardly used in empiric-based dosing, when leukocyte CoQ10 levels were markedly elevated above the reference range (Supplementary Table S1), the family reported subjective clinical improvement coincident with these supraphysiological peaks and perceived decline within months after returning to standard dosing. While anecdotal, this longitudinal observation is consistent with our cellular finding that the mitochondrial phenotype of CS is pharmacologically modifiable. Whether early metabolic intervention can alter long-term disease trajectory in CS warrants prospective investigation in rigorous clinical trials.

Several limitations exist to the preclinical findings identified in CSB-deficient patient fibroblasts, including biochemical variability, autophagic resolution, model systems, and genotypic diversity. In terms of biochemical variability, glutathione endpoint measurements in this cohort were variable across replicates and conditions, suggesting that bulk steady-state GSH/GSSG indices may not fully capture the dynamic oxidative redox constraints revealed by the functional assays. Time-resolved sampling and flux-oriented measures of redox buffering will be important future approaches to align biochemical and phenotypic readouts. For autophagic resolution, LC3 lipidation changes under stress, together with the rapamycin responsiveness and the bafilomycin control, are compatible with altered induction of autophagy/quality control in CSB-deficient cells under the specific stress paradigm used here, but they do not define cargo specificity or the precise step(s) affected. Resolving whether the dominant vulnerability is mitochondrial-targeted clearance versus broader stress adaptation will require future mitophagy-directed assays. Our preclinical model was patient-derived primary fibroblasts that effectively recapitulate the germline genetic defect, but which may not fully capture the extreme bioenergetic demands and specific vulnerability of post-mitotic neurons, which are the primary tissues affected in the clinical neurodegenerative phenotype. Future studies employing induced pluripotent stem cell (iPSC)-derived neurons or cerebral organoids would be valuable to validate these metabolic rescue strategies in a disease-relevant neuronal context. In terms of genotypic diversity, our cohort was restricted to two siblings carrying the same compound heterozygous variants in *ERCC6*, which encodes CSB. Given conflicting reports in the literature regarding CS bioenergetics, these findings need to be validated in a larger Cockayne syndrome cohort representing diverse *ERCC6* genotypes as well as *ERCC8* (CSA) deficiency to determine if these rescue strategies are universal to the Cockayne spectrum.

In summary, fibroblasts from two patients with CS due to compound heterozygous *ERCC6* variants exhibit a stress-sensitive mitochondrial maintenance defect that recapitulates cardinal features of PMD at the cellular level. Mechanistic findings in this study support an overall integrative model in which CSB loss produces a compound mitochondrial maintenance defect — mtDNA depletion, respiratory chain structural compromise, impaired autophagic clearance, and constitutive oxidative redox imbalance — that is partially buffered under nutrient-replete conditions but decompensates when antioxidant capacity and metabolic flexibility are simultaneously constrained. Importantly, pharmacological rescue of CSB-deficient cell viability was achievable across four distinct levels: (i) upstream reduction of mitochondrial oxidant burden primarily through GSH replenishment (NAC) or electron carrier support (CoQ10); (ii) restoration of mitochondrial quality control by bypassing the CSB-dependent autophagy induction defect (rapamycin); (iii) reinforcement of extra-mitochondrial antioxidant defense through transcriptional pathway activation ((−)-epicatechin); and (iv) increased cellular tolerance to the consequences of acute GSH depletion through mitochondrial translation fidelity support and membrane stabilization (taurine). The coexistence of four effective rescue axes demonstrates that CS-associated mitochondrial vulnerability arises from convergent failure of multiple homeostatic systems rather than a single biochemical bottleneck. Thus, pharmacologic treatment of mitochondrial dysfunction in CS via antioxidant supplementation, respiratory chain support, autophagy modulation, and mitochondrial translational maintenance identifies multiple tractable targets for therapeutic development, which may be complementary to strategies addressing the primary DNA repair deficiency. Rigorous clinical trials are warranted in Cockayne syndrome to objectively evaluate the safety and efficacy of these lead therapeutic strategies that mitigate oxidative stress, improve autophagy induction, and improve mitochondrial bioenergetics on patient survival, function, and feeling.

## Supporting information

CSB-LC3-Gel1

CSB-LC3-Gel2

CSB-LC3-BetaAct-Gel1

CSB-LC3-BetaAct-Gel2

SourceData_Figure2

SourceData_Figure4

SourceData_Figure5

SourceData_Figure6

SourceData_Figure3

## Ethics Declaration

This study was approved by the Institutional Review Board at Children’s Hospital of Philadelphia (#08-6177, Falk PI). Written informed consent was obtained from the patients’ parents for participation in the study and for the use of clinical data and biospecimens. The study was conducted in accordance with the Declaration of Helsinki. <u>Consent for publication</u>: Written informed consent for publication of clinical details was obtained from the patients’ parents.

## Acknowledgements.

We are grateful to the patients and their family for inspiring and participating in this study, as well as Nicholas Wachowski, Atif Towheed, PhD, and Jennifer Murray for their assistance and guidance in this effort. This work was funded in part by the National Institutes of Health (R35-GM134863 to MJF and U54-NS078059 NAMDC fellowship to MK) and the philanthropically funded CHOP Mitochondrial Medicine Fellowship Program. The content is solely the responsibility of the authors and does not necessarily represent the official views of the funders, including the NIH.

## Author Contributions

MJF conceived of, designed the study, obtained study funding, provided long-term clinical care for the study subjects, reviewed all study data and figures, and assisted in manuscript preparation and editing. MK performed phenotypic characterization, cell culture, and experimental work, including cell survival assays, mitochondrial stress assessment (MitoSOX, CX5 CellInsight), cellular oxidative stress assessment (CellROX, flow cytometry), western immunoblots and data analysis. EMM oversaw IRB review and collected and curated the clinical data on the study subjects. SH provided mentorship on experimental design and manuscript preparation. CR helped to optimize the CX5 CellInsight high content image protocol, prepared the positive control plate, helped to design and analyze the autophagy marker western blot experiments. KK performed the analysis of the positive control compound screen. ENO performed the GSSG/GSH analyses and provided mentorship on experimental design. MK drafted the manuscript. All authors reviewed and approved the final manuscript.

---

**Supplementary Figure S1 | NAC rescues metabolic stress-induced ATP collapse in CSB-deficient fibroblasts when administered concurrently or after stress onset.**

**(a)** ATP-based cell survival (CellTiter-Glo luminescence, expressed as percentage of basal DMEM) in control (open circles) and CSB-deficient Patient 1 (open diamonds) fibroblasts under basal DMEM, galactose-only stress (galactose 10 mM, L-glutamine 1 mM), and combined stress with concurrent NAC administration (galactose 10 mM, L-glutamine 0.5 mM, BSO 50 µM, NAC 2 mM). In the concurrent protocol, BSO and NAC were added simultaneously at 24 hours after cell seeding; luminescence was measured at 72 hours. Schematic (right) illustrates the experimental timeline.

**(b)** Representative phase-contrast micrographs of CSB-deficient (Patient 1) fibroblasts under combined stress conditions (galactose 10 mM, L-glutamine 0.5 mM, BSO 50 µM). Upper row: cells imaged at 24, 48, and 72 hours under stress prior to NAC addition, showing progressive morphological deterioration. Lower row: cells imaged at 96, 120, and 144 hours and at day 10 following addition of NAC 2 mM at 72 hours (i.e., 7 days of NAC treatment), demonstrating partial morphological recovery.

**(c)** Post-stress rescue paradigm. Patient-1 and CTRL fibroblasts were exposed to stress for 48 h prior to addition of 2 mM NAC; ATP readout was performed 48 h after NAC addition. Right: treatment timeline.

Total intracellular ATP was quantified by CellTiter-Glo luminescence assay and expressed as percent of CTRL under basal DMEM. Bars represent mean ± s.d.; individual data points represent independent biological replicates. Statistical comparison by one-way ANOVA with Tukey’s post hoc test. \*\*\*\**P* < 0.0001. Data are expressed as mean ± s.d.; *n* = 3 independent biological replicates. Statistical comparison by one-way ANOVA. ****P < 0.0001. BSO, buthionine sulfoximine; NAC, N-acetylcysteine.

**Supplementary Figure S2 | Representative CellROX Deep Red flow cytometry histogram plots for five dual-rescue compounds.**

Representative flow cytometry histograms depicting total cellular ROS levels in control and CSB-deficient Patient 1 fibroblasts across all experimental conditions, shown for each of the five dual-rescue compounds: CoQ10, taurine, (−)-epicatechin, and rapamycin (one representative replicate per compound shown; *n* = 3 independent biological replicates per condition). CellROX Deep Red fluorescence intensity is plotted on the x-axis (R1-A, area parameter) and cell count on the y-axis. Conditions shown per panel, from top to bottom: unstained control (autofluorescence baseline); positive control (TBHP-treated, defining the maximal inducible oxidative shift); Control DMEM; Patient DMEM; Control DMEM + compound; Patient DMEM + compound; Control Galactose; Patient Galactose; Control Galactose + compound; Patient Galactose + compound; Control Galactose + BSO; Patient Galactose + BSO; Control Galactose + BSO + compound; Patient Galactose + BSO + compound. A rightward shift in fluorescence intensity relative to the Control DMEM condition indicates increased cellular ROS burden. Quantification of median fluorescence intensity (MFI) across all three biological replicates for each condition and compound is presented in Fig. 5d–h. BSO, buthionine sulfoximine; TBHP, tert-butyl hydroperoxide.

**Supplementary Figure S3 | Uncropped western blot images for Figure 6**.

Raw, uncropped immunoblot membranes corresponding to the quantified data presented in Fig. 6. Upper left panel: p62 (SQSTM1), LC3-II, and LC3-I bands in control (C) and CSB-deficient Patient 1 (P) fibroblasts under basal DMEM, galactose + BSO (combined stress), galactose + BSO + rapamycin, and galactose + BSO + bafilomycin A1 conditions. Upper right panel: continuation of the same membrane showing galactose + BSO + bafilomycin A1 + rapamycin, galactose-only, and basal DMEM control conditions. Molecular weight markers are visible on the left of each panel; band identities (p62, LC3-II, LC3-I) are indicated by red arrows. Lower panels: uncropped β-actin and TOM20 loading control membranes across the corresponding conditions and lane arrangements. All membranes were processed and imaged as described in the Methods. Regions used for quantification in Fig. 6b–c are indicated by the cropped areas shown in the main figure. C, control fibroblasts; P, CSB-deficient Patient 1 fibroblasts; BSO, buthionine sulfoximine; Gal, galactose; Rap, rapamycin; Baf A1, bafilomycin A1.

**Supplementary Table S1.** Detailed clinical features of CS Patients 1 and 2.

**Supplementary Table S2.** Biochemical and histopathological parameters CS Patients 1 and 2.

