## Supplementary Table for "Targeting the Mitochondrial Phenotype in Cockayne Syndrome Patient Cells: From Bioenergetic Fragility to Pharmacologic Rescue"

| Supplementary Table 1: Summary of clinical biochemistry and corresponding treatment dosing of two siblings | | | | | |
| --- | --- | --- | --- | --- | --- |
|  | | Patient 1 | | Patient 2 | |
|  | | Value | Age (y) | Value | Age |
| Blood Lactate (mmol/L) | | 1.8-2.73 | 4-17 | 1.6-2.2 | 2-14 |
| Blood Pyruvate (mmol/L) | | 0.1-0.18 |  | 0.12-0.16 |  |
| Anion Gap (mEq/L) | | 10-20 |  | 6-13 |  |
| Blood Alanine  *(Ref: 186-524 µmol/L)* | | 215.7-639.5 |  | 251.8-603.8 |  |
| Acylcarnitine profile | | Normal |  | Normal |  |
| Glutathione Profile and NAC administration | | | | | |
| Glutathione Profile and NAC administration | NAC dosing | 10 mg/kg/d | 8-10 | 10mg/kg/d | 6-8.5 |
|  |  | 15 mg/kg/d | 10-12 | 18mg/kg/d | 10-13 |
|  |  | 20 mg/kg/d | 12-18 | 30 mg/kg/d | 13-16 |
|  | Total  *(Ref:544-1228 µmol/L)* | 199-856 | 10-18 | 314-1154.4 | 8-16 |
|  | Reduced  *(Ref:544-1228 µmol/L)* | 751.7-855.9 | 16-18 | 1073-1152.5 | 14-16 |
|  | Oxidized  *(Ref: <1.78 µmol/L)* | 0.88 -1.04 |  | 1.09-1.85 |  |
|  | Reduced/Oxidized  *(Ref: 597-2584))* | 476.4 - 823 |  | 622.7-984.4 |  |

| Leukocyte CoQ10 Profile and ubiquinol administration | | | | | |
| --- | --- | --- | --- | --- | --- |
| Patient 1 | | | Patient 2 | | |
| Age (years) | Leukocyte CoQ10  **: (MNG Lab - Ref:66-183 pmol/mg/protein)*  ***: (ARUP -Ref: 0.4-1.6 mg/L)* | Ubiquinol administration  (mg/kg/d) | Age (years) | Leukocyte CoQ10  **: (MNG Lab - Ref:66-183 pmol/mg/protein)*  ***: (ARUP -Ref: 0.4-1.6 mg/L* | Ubiquinol administration  (mg/kg/d) |
| 6-7 | 174-241 (*) | 13-14 | 4-5 | 219-238 (*) | 12-13 |
| 8-10 | 253- 458 (*) | 140-300 | 6-8 | 263-693 (*) | 70-300 |
| 10-13 | 204-139 (*) | 8-16 | 8-13 | 240-184 (*) | 8-10 |
| 13-17 | 2.9-3.6 mg/dl (**) | 30 | 13-16 | 3.2 - 4.17 (**) | 30 |

| Supplementary Table S2: Summary of diagnostic workup and mitochondrial biochemistry in two siblings. | | | | | |
| --- | --- | --- | --- | --- | --- |
|  | | Patient 1 | | Patient 2 | |
| Fragile X analysis (LabCorp) | | (3y) Normal CGG repeat range (<45) | | *N/A* | |
| SNP Microarray | | (4y) No clinically significant CNVs (46,XY) | | *N/A* | |
| Vanishing white matter panel (*EIF2B1, EIF2B2, EIF2B3, EIF2B4*, and *EIF2B5)* | | (4y) No pathogenic variant | | *N/A* | |
| *PLP1* deletion and duplication analysis (Greenwood) | | (4y) No pathogenic variant | | *N/A* | |
| *PLP1* sequencing, deletion/ duplication analysis | | (4y) No pathogenic variant, no deletion-duplication | | *N/A* | |
| *GJC2* gene sequencing (Baylor) | | (4y) No pathogenic variant | | (2y) No pathogenic variant | |
| Mitochondrial Disease Panel (101 nuclear PMD causing genes + mtDNA sequencing (GeneDx) | | (4y) No pathogenic variant | | (2y) No pathogenic variant | |
| X-linked ID Panel | | (4y) No pathogenic variant | | *N/A* | |
| *GALC* gene sequencing, deletion-duplication (Emory) | | (6y) No pathogenic variant, no deletion-duplication | | (4y) No pathogenic variant, no deletion - duplication | |
| Leukodystrophy NGS panel (*ABAT, ACOX1, ALDH3A2, ARSA, ASPA, DARS2, EIF2B1, EIF2B2, EIF2B3, EIF2B4, EIF2B5, FAM126A, GFAP, GJC2, HEPACAM, HSPD1, HTRA1, MLC1, NOTCH3, PLP1, PSAP, PTEN, SCP2, SLC25A12, CSF1R*, and *LMN81*) (Fulgent Diagnostics) | | (6y) No pathogenic variant | | *N/A* | |
| Whole exome sequencing (Emory) | | (4y) *SPG11:*c.4070T>C (Paternal) | | (2y) *SPG11*:c.4070T>C (Paternal) | |
| Urine oligosaccharide & glycan analysis | | (7y) Abnormal peak  *m/z = 1449* | | (5y) Abnormal peak  *m/z = 1449* | |
| Whole exome sequencing *reanalysis* (Emory) | | (7y) *ERCC6:* c.1526+1G>T / c.2800C>A; p.(P934T) | | (5y) *ERCC6: c*.1526+1G>T / c.2800C>A; p.(P934T) | |
| Muscle  Biopsy | Muscle histopathology (CHOP) | Minimal, non-specific changes; no morphological evidence of mitochondrial myopathy | |  | |
|  | Muscle CoQ10 (MNG Labs) | 238 pmol/mg | |  | |
|  | mtDNA content (Baylor) | 38% of control (consistent with mtDNA depletion) | | N/A | |
|  | mtDNA sequencing (Baylor) | m.8821T>C variant 🡪 3.8% heteroplasmy. | |  |  |
|  | Complex I | 156% | |  |  |
|  | Complex I+III — total | 112% | |  |  |
|  | Complex I+III — rotenone-sensitive | 131% | |  |  |
|  | Complex II | 132% | |  |  |
|  | Complex II+III | 115% | |  |  |
|  | Complex IV | 148% | |  |  |
|  | Citrate synthase | 123% | |  |  |
| Skin Biopsy  Fibroblast respiratory chain enzyme activities  (Baylor) |  | Before correction for CS | After correction for CS | Before correction for CS | After correction for CS |
|  | Complex I | 126% | 85 (%) | 118% | 178 (%) |
|  | Complex I+III — total | 167% | 113 (%) | 133% | 200 (%) |
|  | Complex I+III — rotenone-sensitive | 556% | 375 (%) | 194% | 293 (%) |
|  | Complex II | 181% | 122 (%) | 94% | 141 (%) |
|  | Complex II+III | 210% | 141 (%) | 92% | 139 (%) |
|  | Complex IV | 229% | 154 (%) | 82% | 123 (%) |
|  | Citrate synthase | 148% | | 66% | |

**Abbreviations:** CNV, copy number variant; CS, citrate synthase; ID, intellectual disability; mtDNA, mitochondrial DNA; PMD: Primary mitochondrial disorder; SNHL, sensorineural hearing loss; CoQ10, coenzyme Q10; N/A, not applicable.
