## Supplementary figures and images for "Targeting the Mitochondrial Phenotype in Cockayne Syndrome Patient Cells: From Bioenergetic Fragility to Pharmacologic Rescue"

### CSB-LC3-BetaAct-Gel1

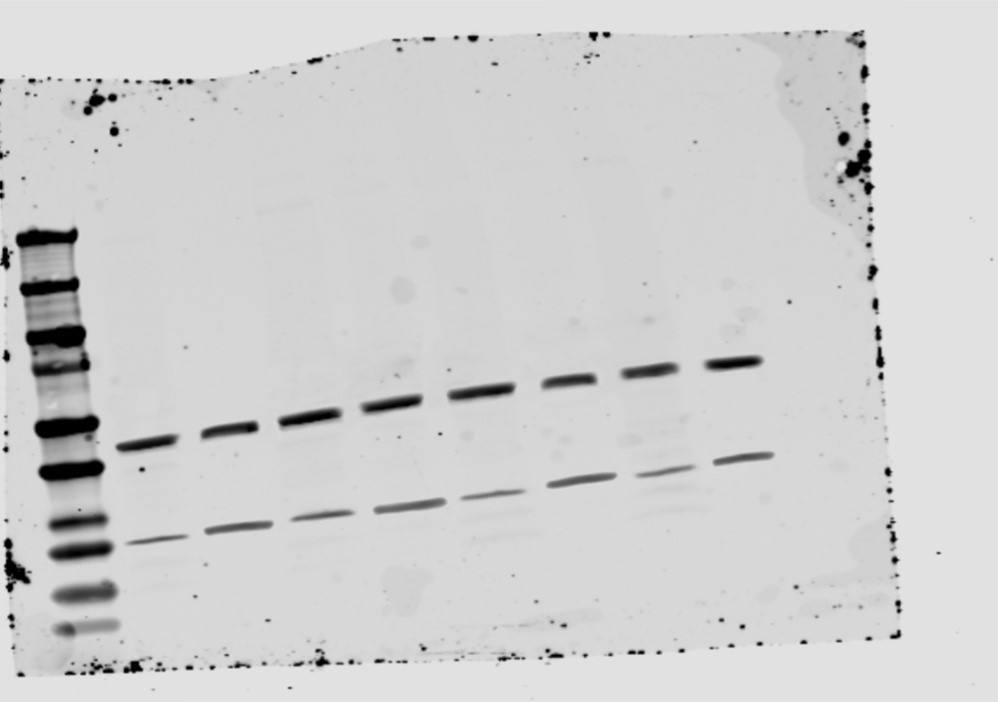

### CSB-LC3-BetaAct-Gel2

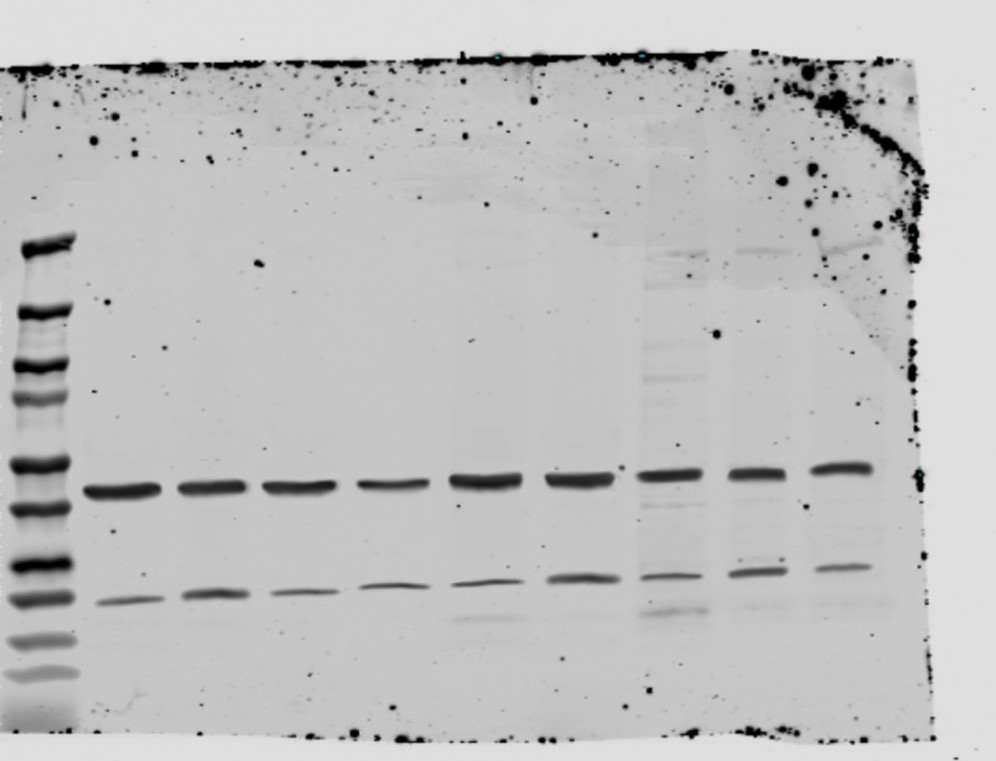

### CSB-LC3-Gel1

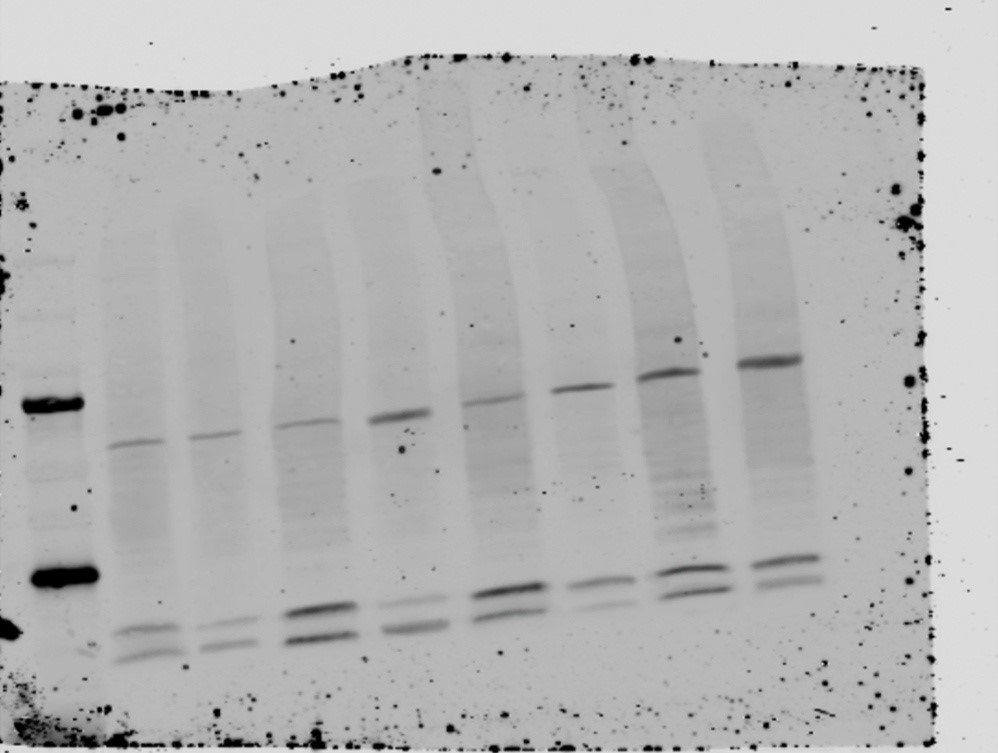

### CSB-LC3-Gel2

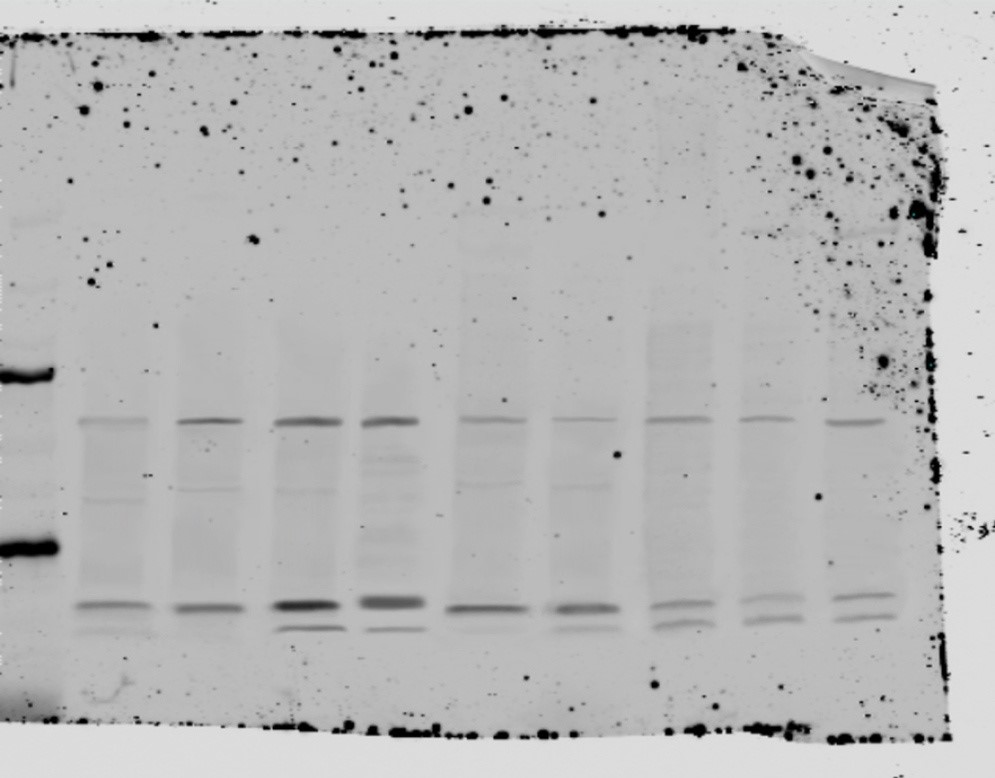

### Supplemental Figure

#### Figure S1.

A

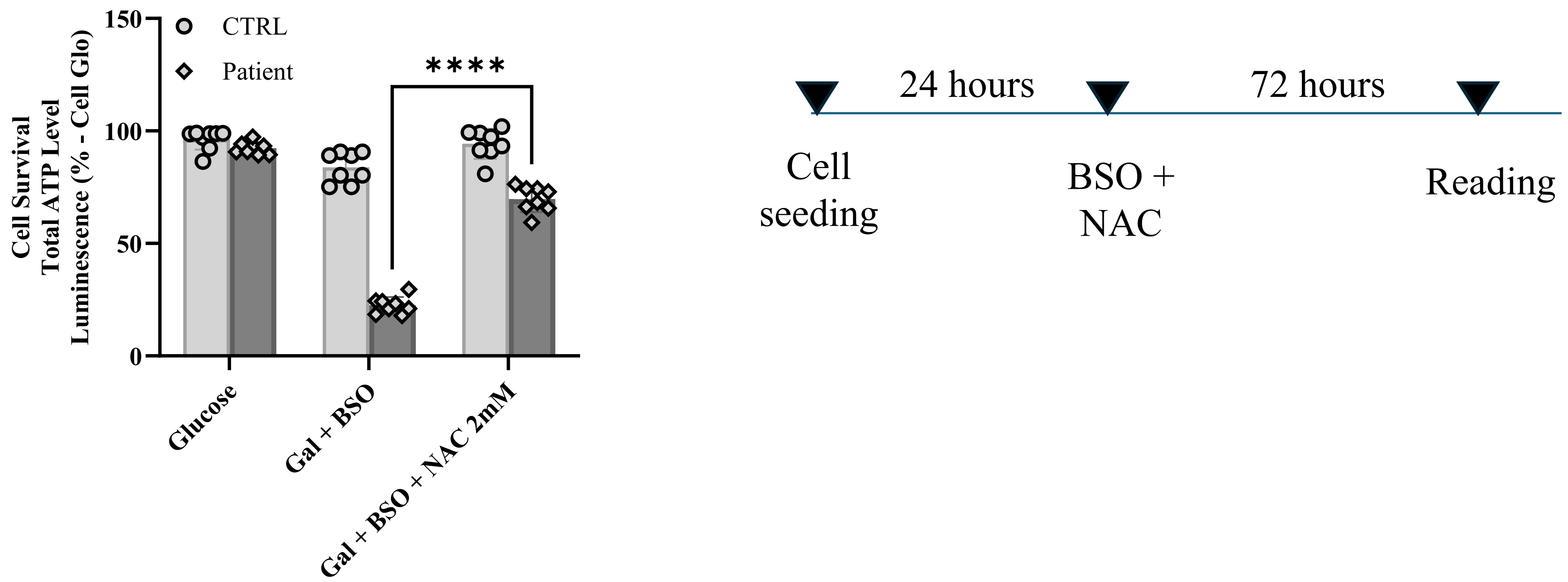


B

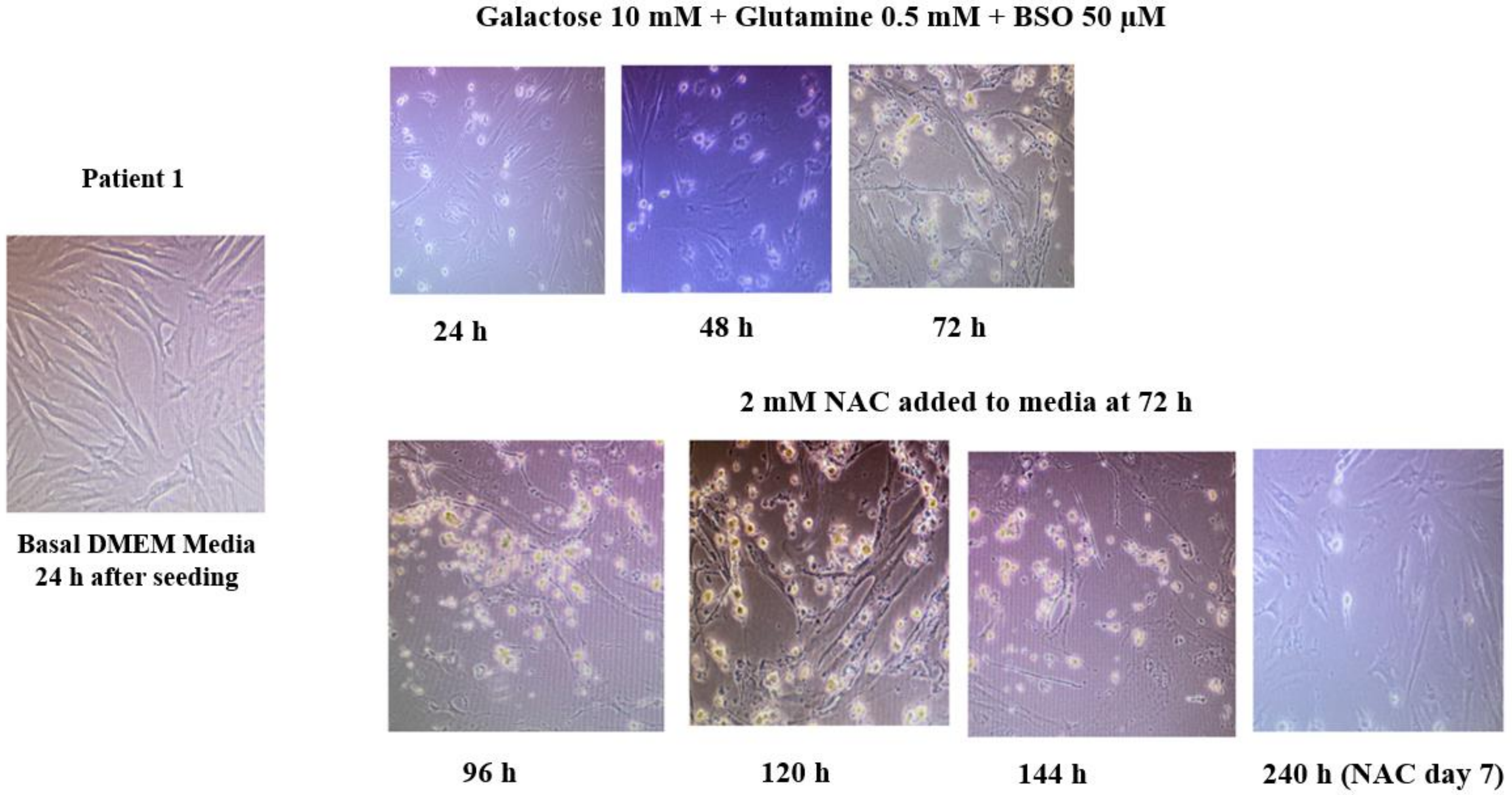


C

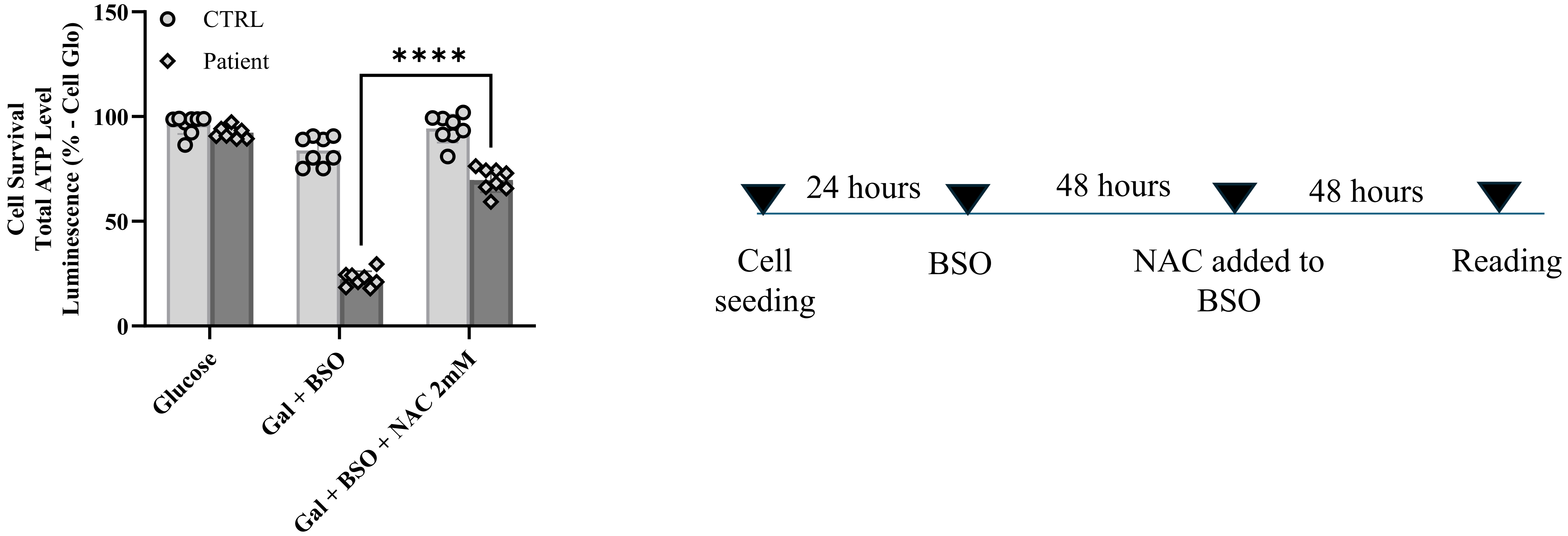

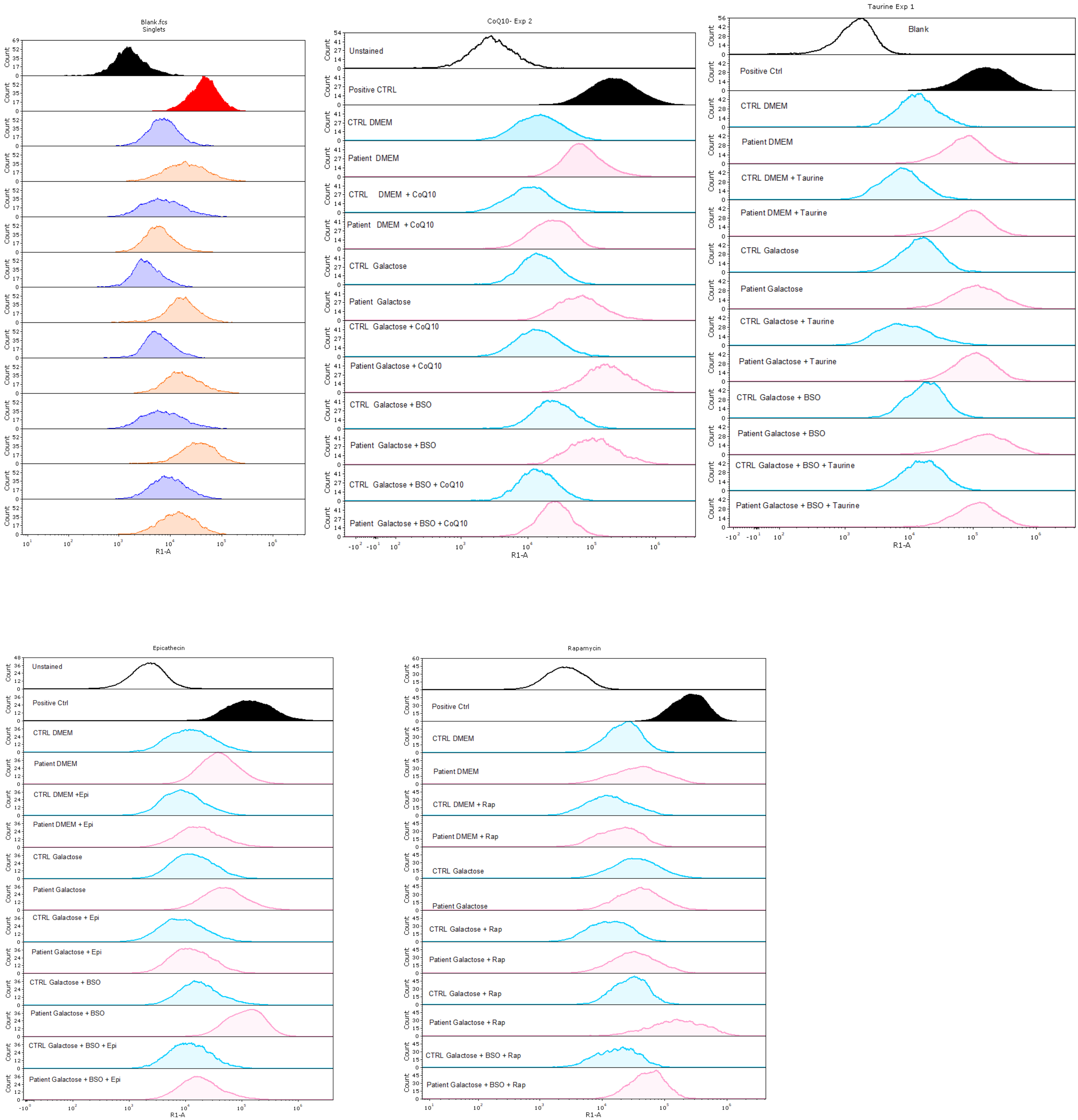

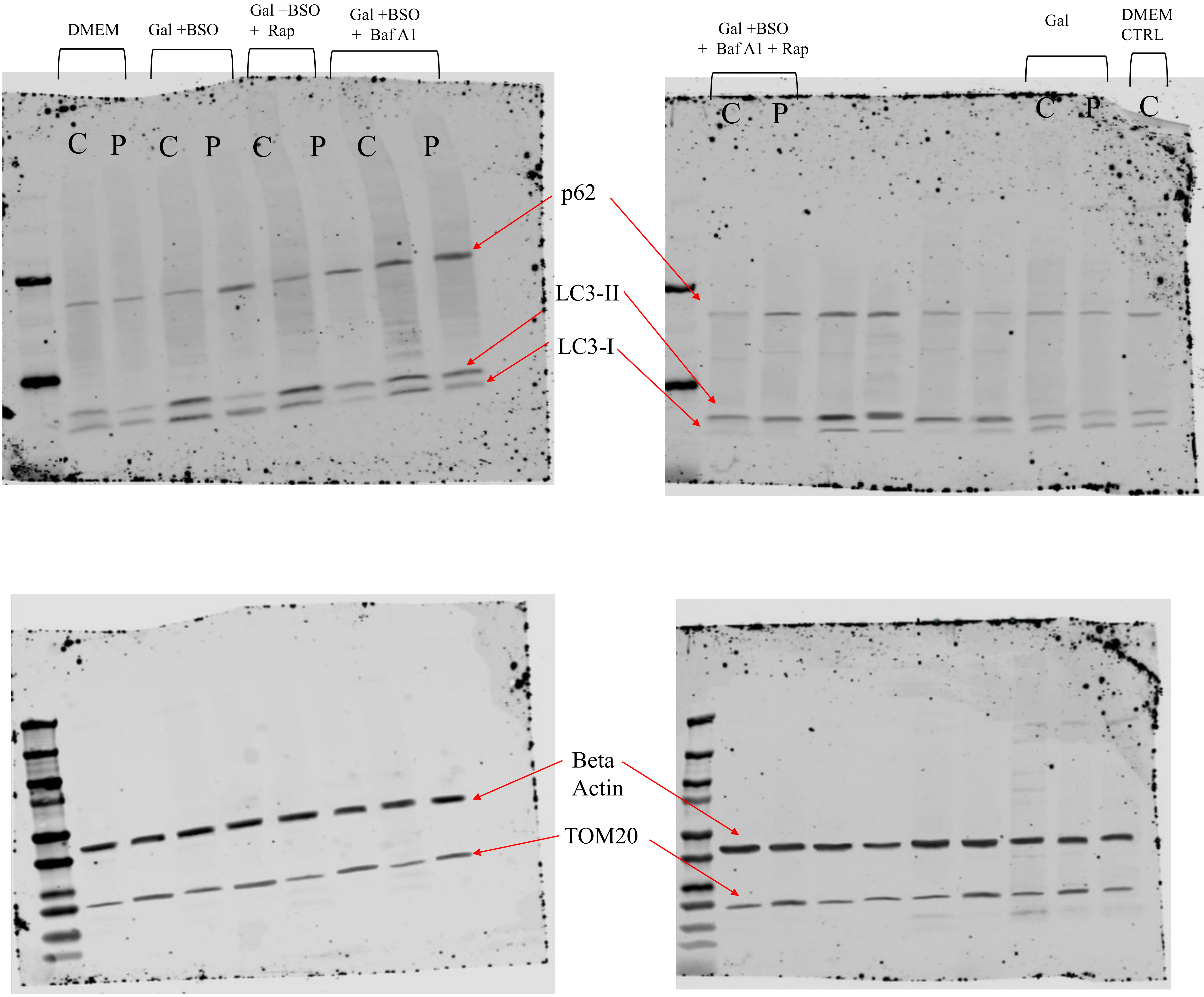
